# Sequential narrative binding by hippocampal CA2/3 sustains lifespan episodic detail retrieval

**DOI:** 10.64898/2026.03.17.712337

**Authors:** Thomas D. Miller, Joseph H. Zhou, Adam E Handel, Thomas A. Pollak, Penny A. Gowland, Chrystalina A. Antoniades, Michael S. Zandi, Clive R. Rosenthal

## Abstract

Hippocampal subfield contributions to the structure of episodic memories across the lifespan remain incompletely understood, particularly regarding the mechanisms underlying sustained retrieval of detailed events regardless of age. Here, we show that focal hippocampal damage in 32 individuals with amnesia produces time-invariant loss of episodic content details (event specifics, place, perceptual, and emotional content) across six decades, sparing only early childhood memories and semantic details. Temporal context details showed non-monotonic vulnerability, with maximal deficits in recent memories. Volumetric analyses revealed distinct anterior-posterior functional gradients: anterior hippocampal CA3 predicted retrieval across 11–55+ years, posterior CA3 across 18–55+ years, whereas CA1 predicted only recent memories. Computational linguistic analyses using Sentence-BERT semantic embeddings quantified narrative organisation within remembered events via consecutive-sentence semantic relatedness (local narrative coherence) and overall semantic alignment (global narrative coherence). Hippocampal CA2/3 damage impaired local but not global narrative coherence, with reduced local coherence mediating episodic detail loss. This dissociation demonstrates that CA2/3 mediates sequential narrative binding through mechanisms dissociable from broader narrative operations. Concordant time-invariant deficits across quantitative measures of episodic content and computational-linguistic based measures challenge classical systems consolidation predictions, supporting parallel hippocampal-cortical contributions to lifespan episodic memory.

## Introduction

Human autobiographical episodic memories are first-person, re-experiential events that depend on dynamic interactions between the hippocampus and distributed neocortical networks ^1–5^. Lesion studies in humans provide conflicting evidence about the duration of hippocampal involvement in memory consolidation and retrieval, with some studies reporting time-limited hippocampal contributions ^6–8^, whereas others find persistent deficits for detailed lifespan memories ^9–11^. In parallel, functional neuroimaging studies find graded hippocampal involvement with memory remoteness ^12,13^ and memory age–independent activity for vivid recall ^2^.

Many of these prior investigations treated the hippocampus as a single, undifferentiated structure. However, the hippocampus exhibits functional specialisation along its long axis, with anterior regions preferentially supporting schematic, gist-level retrieval, whereas posterior regions exhibit a temporal gradient that favours recent, detail-rich memories ^14–22^. Hippocampal subregional specialisation is embedded within this functional neuroanatomic organisation ^15,23^, revealing separable functional contributions of individual subfields ^24–28^.

Three key empirical gaps remain. First, how anterior–posterior specialisation manifests when retrieving memories from across the lifespan remains incompletely characterised. Although several fMRI investigations have reported anterior–posterior gradients in hippocampal activity for recent versus remote events ^14,15^, few studies have examined whether structural hippocampal subfield measures, particularly in human focal lesion cohorts, track these functional gradients. Second, the content-specific vulnerability of episodic components (temporal, spatial, perceptual, emotional) across lifespan consolidation lacks systematic investigation in causal human lesion cohorts. Content-specific vulnerability may reflect specialised processing across distinct hippocampal subfields that contribute to the encoding and maintenance of specific episodic features ^6,24,25^. Third, how hippocampal subfields contribute to the structural integration of narrative elements into coherent autobiographical memories has not been quantified in the context of hippocampal amnesia.

In the current study, hippocampal subfield contributions to the retrieval and integration of narrative elements into coherent autobiographical memories from across the lifespan were investigated in a human model of focal lesion damage. Leucine-rich glioma inactivated 1-limbic encephalitis (LGI1-LE) provides a foundational model to test causal contributions of CA2/3 and CA1 to episodic consolidation and retrieval, with minimal extra-hippocampal pathology ^11,29,30^. Data from 32 individuals with CA2/3 and CA1 damage secondary to LGI1-LE and 32 matched controls were analysed using three complementary approaches that tested specific computational and behavioural predictions.

First, the Autobiographical Interview ^31^, an established quantitative measurement tool, was used to examine memory consolidation across the lifespan. We hypothesised that CA2/3 volume would predict episodic retrieval performance across six decades of adult life, reflecting its role in rapid high-fidelity item–context binding irrespective of memory age, whereas early-childhood events and semantic details would remain spared ^11,32–34^. Conversely, CA1 was predicted to impair only recent episodic retrieval, because temporal interference from newer events places greater demands on CA1-mediated pattern separation ^35,36^. Second, we examined content-specific deficits by parsing memories into context-bound components: event details, temporal information, spatial location, perceptual content, and emotions/thoughts. CA2/3 damage was hypothesised to disrupt all of these components across the lifespan, though temporal details, providing explicit chronological scaffolding for remembered events ^37–40^, may vary over lifespan retention intervals given distinct computational demands of recent versus remote representations ^2,41^. Third, event structure was assessed using SBERT-based semantic embeddings to compute local narrative coherence (cosine similarity between adjacent sentences) and global narrative coherence (mean similarity between sentences and the narrative centroid) ^42,43^. These constructs extend computational discourse coherence frameworks, previously applied to cognitive aging and dementia ^44–46^, to hippocampal amnesia. Hippocampal CA2/3 damage was hypothesised to impair local coherence whilst preserving global narrative coherence, reflecting how the recurrent collateral architecture of CA3 mediates rapid binding of adjacent episodic elements within local contexts ^47,48^ versus distributed hippocampal–cortical networks supporting schema integration ^49–53^.

Our findings reveal three novel empirical patterns. First, hippocampal CA2/3 damage yielded time-invariant loss of episodic detail across six decades, whereas CA1 volume predicted only recent episodic details, revealed through distinct anterior–posterior gradients: anterior CA3 from early adulthood (age 11) through older age (55+), posterior CA3 from ages 18–55+, and anterior/posterior CA1 solely for recent events. Second, all content components were lifespan-impaired and predicted by CA3 volume, with non-monotonic temporal detail vulnerability (greatest CA2/3-related impairment for recent memories and early-adulthood memories versus minimal for ages 18–30), whereas CA1 predicted only recent event details. Third, local narrative coherence was disrupted across the lifespan by CA2/3 damage, whereas global narrative coherence remained intact; mediation analysis showed that impaired local coherence accounted for the relationship between CA2/3 damage and episodic amnesia. The convergence of time-invariant deficits across quantitative episodic content measures and computational linguistic analyses reveals distinct computational contributions of hippocampal subfields to lifespan episodic memories: CA3 mediates sequential binding of episodic elements through local narrative coherence, whereas CA1 contributes selectively to recent memory retrieval.

## Results

### Participant characteristics and anatomical findings

Results from 18 individuals with amnesia secondary to LGI1-LE revealed focal bilateral hippocampal pathology localised at 3.0-Tesla to CA1 and CA2/3 ^30^, consistent with focal bilateral CA3 damage localised at 7.0-Tesla in our cohort of 14 individuals ^11^. Both cohorts exhibited greater CA2/3 than CA1 atrophy (see Supplementary data 1) ^11,29,54^. Voxel-based morphometry revealed no suprathreshold grey matter volume loss in either cohort relative to controls. Despite differing MRI acquisition parameters and magnetic field strengths, both cohorts showed consistent VBM results matching reports from other laboratories on chronic LGI1-LE pathology ^11,29,55–62^. Comprehensive neuropsychological assessment (Supplementary Table S1) revealed isolated impairment in verbal memory performance alongside preserved or supranormal performance across multiple cognitive domains, including language, executive function, semantic processing, and visuospatial abilities. Crucially, preserved language and semantic processing ensure that observed deficits in episodic detail retrieval and computational linguistic measures of narrative coherence (derived from sentence-level semantic embeddings) reflect hippocampal-specific memory dysfunction rather than language or general cognitive impairment. The neuropsychological profile establishes LGI1-LE as a selective model of hippocampal amnesia, distinct from aetiologies with broader cognitive or neuroanatomical compromise. Figure 1 illustrates the anatomy and segmentation protocols used to determine anterior and posterior hippocampal subfield volumes.

**Figure 1.**
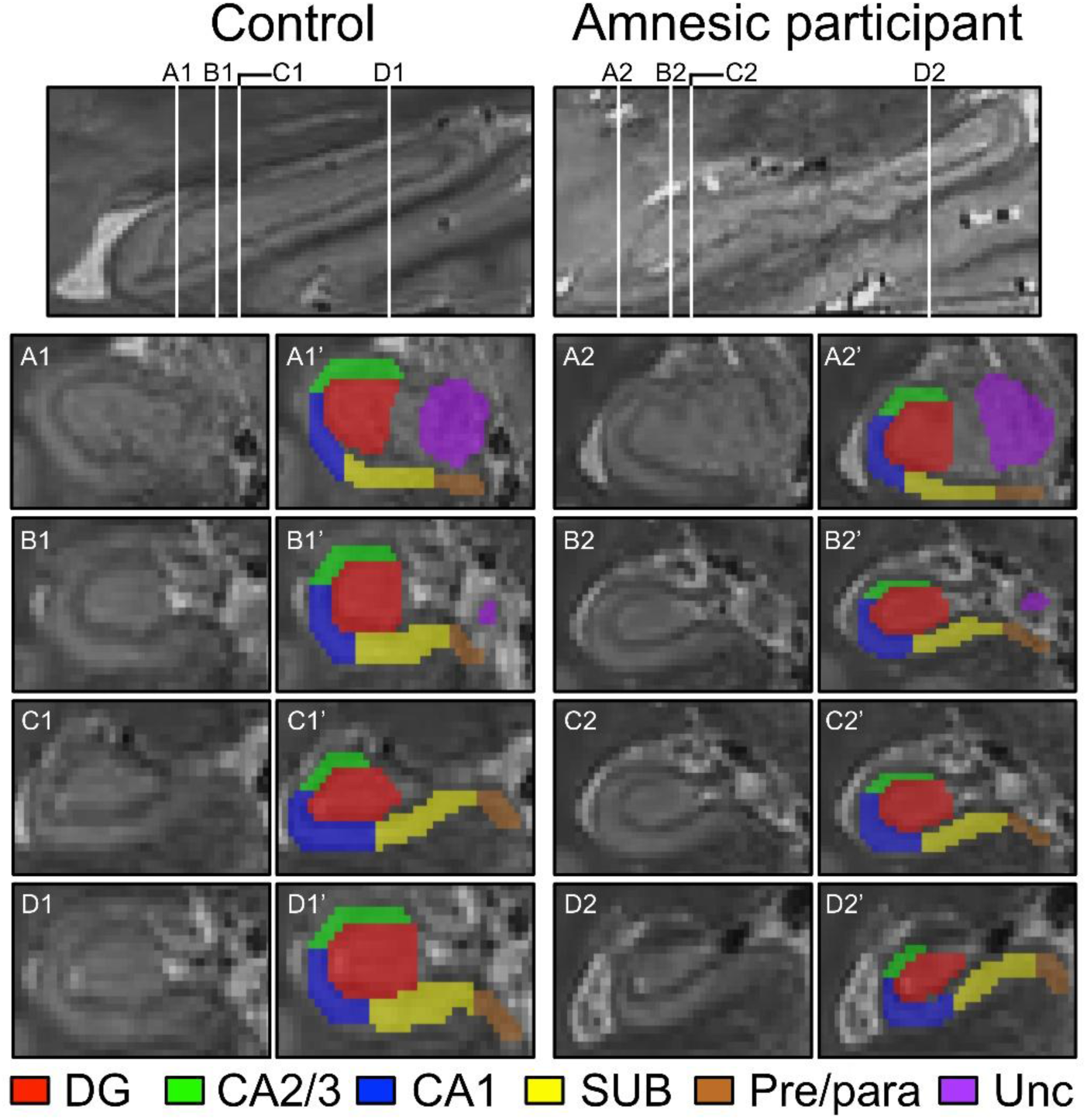
Quantitative T2-weighted 3.0-Tesla high-resolution hippocampal subfield imaging. Sagittal images (top row) illustrate the full longitudinal axis of the hippocampus in a representative control participant and participant with hippocampal amnesia acquired using 3.0-Tesla MRI. Coronal images (rows A–D) display colour-coded hippocampal subfield segmentation results in a control and participant with amnesia (denoted as,’). The anterior-posterior boundary is demarcated at the transition between panels B and C. Abbreviations: CA1, cornu Ammonis 1; CA2/3, cornu Ammonis 2 & 3; DG, dentate gyrus; Pre/para, pre/parasubiculum; SUB, subiculum; Unc, uncus.

To enable cross-cohort structure-function analyses despite differing MRI acquisition parameters, hippocampal subfield volumes were harmonised across datasets using complementary segmentation approaches. For the 7.0-Tesla cohort, composite CA2/3 volume was calculated as the sum of CA2 and CA3 volumes, reflecting their functional integration as a subregion ^63,64^. Z-scores for total CA2/3 volume were then computed across both cohorts, enabling standardised comparison despite different acquisition protocols ^65,66^. For the 3.0-Tesla cohort, where lower spatial resolution precluded reliable CA2/CA3 delineation, empirically derived proportional contributions of CA3 and CA2 to composite CA2/3 volumes were estimated from the 7.0-Tesla reference dataset, with separate anterior and posterior ratios applied to accommodate regional heterogeneity ^14,63^. These harmonised CA2/3 volume estimates, alongside CA1 volumes from both cohorts, were used as independent variables in subsequent structure-function analyses ^67^.

### Hippocampal damage leads to time-invariant episodic amnesia

The new, independent LGI1-LE cohort characterised at 3.0-Tesla (N = 36: 18 with amnesia and 18 matched controls) replicated findings from the original 7.0-Tesla cohort, revealing extensive anterograde and retrograde amnesia on the Autobiographical Interview. A mixed-model ANOVA on total detail scores 2 (group: amnesic, control) by 2 (detail type: internal, external) by 5 (time period: last year; 30–55, 18–30, 11–18, 0–11 years) revealed significant main effects of group (*F*_(1,34)_ = 15.70, *p* < 0.001, η² = 0.32), time period (*F*_(3.54,122.11)_ = 10.56, *p* < 0.001, η² = 0.24), and detail type (*F*_(1,34)_ = 400.11, *p* < 0.001, η² = 0.92), with significant group by time period (*F*_(1,34)_ = 11.69, *p* = 0.002, η² = 0.26) and group by detail type (*F*_(1,34)_ = 11.69, *p* = 0.002, η² = 0.26) interactions but no three-way interaction (Figure 2A–B; see Supplementary data 2 for full results and degrees of freedom, df, correction). Planned comparisons revealed no significant group differences for the earliest childhood period (0–11 years) for internal (*F_(_*_1,34)_ = 0.50, *p* = 0.48, η² = 0.02) or external details (*F*_(1,34)_ = 2.54, *p* = 0.12, η² = 0.04). Excluding this interval, a follow-up ANOVA confirmed a robust group by detail type interaction (*F*_(1,34)_ = 13.69, *p* < 0.001, η² = 0.29), driven by significantly reduced internal details (*F*_(1,34)_ = 17.35, *p* < 0.001, η² = 0.34) but preserved external details (*F*_(1,34)_ = 1.32, *p* = 0.26, η² = 0.04). This indicates selective episodic impairment across post early-childhood periods. These results for the standard administration of the Autobiographical Interview were replicated when the 3.0- and 7.0-Tesla cohorts were combined (n = 32 group with amnesia and n = 32 controls, see Supplemental data 2; *N* = 64).

**Figure 2.**
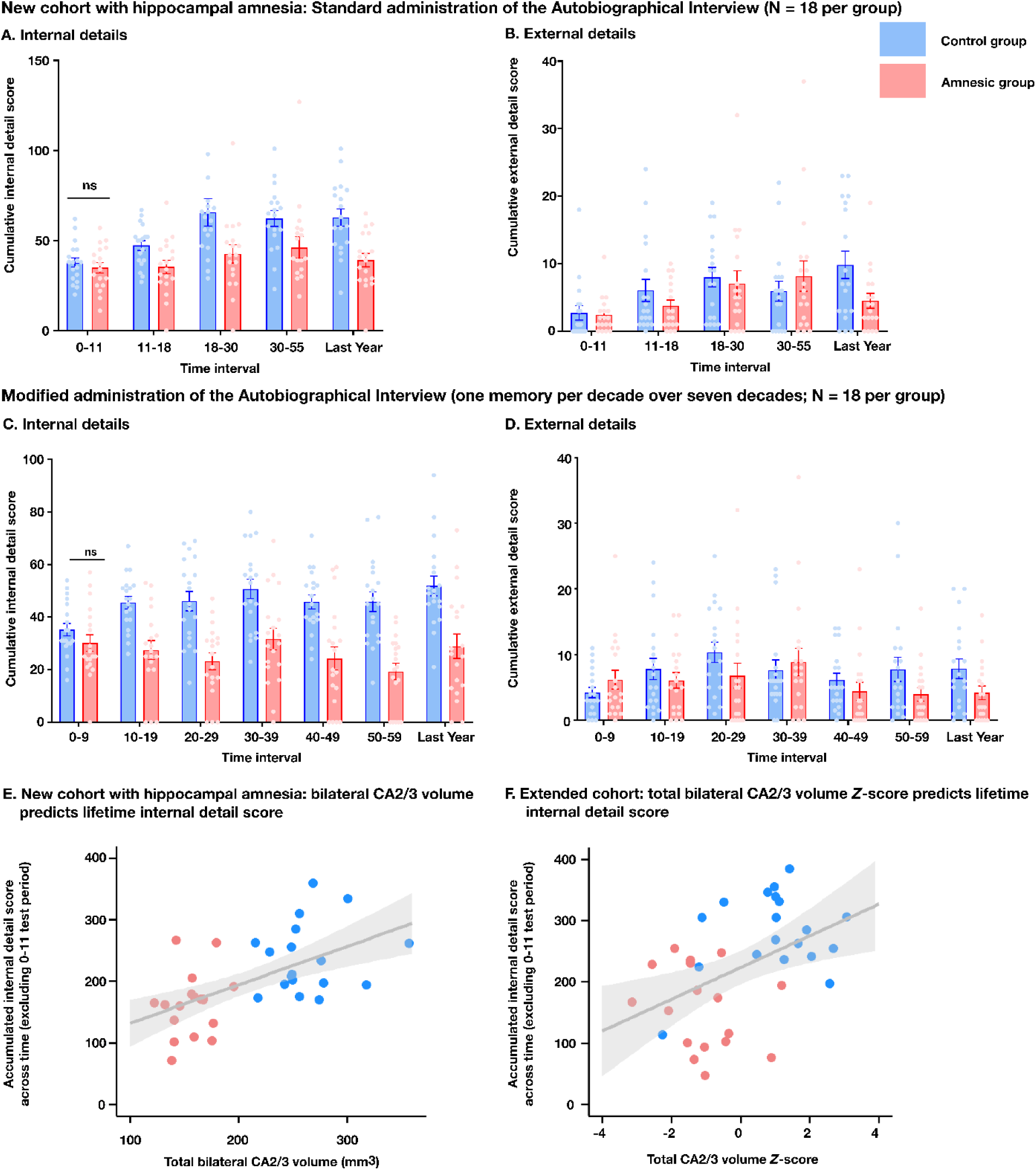
Time-invariant loss of internal (episodic) detail across the lifespan in hippocampal amnesia. (A) Internal (episodic) detail scores across five life periods (0–11, 11–18, 18–30, 30–55 years, and last year) in a new (3.0-Tesla) cohort of participants with hippocampal amnesia (n = 18) and matched controls (n = 18) using the standard Autobiographical Interview. The group with hippocampal amnesia showed significantly reduced internal detail across all post-early-childhood periods (11–18 years onward), with preserved performance for early childhood (0–11 years). (B) External (semantic) detail scores for the same participants across life periods. No significant group differences were observed, demonstrating content specificity of the episodic memory impairment. (C) Internal (episodic) detail scores using the extended decade-by-decade Autobiographical Interview administration in a subset integrated (3.0- and 7.0-Tesla) cohort (n=18) and matched controls (n=18 per group; N=36) across seven time periods (0–9, 10–19, 20–29, 30–39, 40–49, 50–59 years, and last year). The pattern replicates findings from panel A, with time-invariant impairment across post-early-childhood decades and preserved early childhood memory. (D) External (semantic) detail scores for the integrated cohort across the seven time periods. No significant group differences were observed, replicating the content specificity shown in panel B. Full statistical analyses are reported in the main text and Supplementary Tables. (E) Total CA2/3 volume (independent variable) predicts accumulated internal (episodic) detail score (dependent variable, excluding 0–11 years) in the new (3.0-Tesla) cohort (N=36). Stepwise linear regression revealed a significant selective association between CA2/3 volume and episodic detail retrieval (*F*_(1,34)_ = 17.346, *p* < 0.001, R² = 0.338), with CA1 volume not contributing significantly to the model. Each dot represents a single participant (red, amnesia group; blue, control group), with the best-fitting linear regression line shown. (F) Total CA2/3 volume Z-score predicts accumulated internal (episodic) detail score in the subset integrated (3.0- and 7.0-Tesla) cohort using the extended Autobiographical Interview administration (excluding 0–11 years) (N=36). The selective CA2/3 association remains significant (*F_(_*_1,34)_ = 9.031, *p* = 0.005, R² = 0.210), replicating the structure-function relationship shown in panel E. Finally, the full integrated cohort and controls (N = 64: 32 with amnesia (3.0-Tesla cohort, n = 18; 7.0-Tesla cohort, n = 14; 32 matched controls) revealed a significant model (*F*_(1,62)_ = 14.570, *p* < 0.001), with CA2/3 Z-score predicting total interval details (excluding 0-11 years) (*t* = 3.817, p<0.001 R² = 0.190, β₁ = 0.436), whereas the CA1 Z-score was excluded from the model (*t* = -0.435 *p* = 0.665). Each dot represents a single participant (red, amnesia group; blue, control group), with the best-fitting linear regression line shown. Full statistical analyses are reported in the main text and Supplementary Tables. Error bars represent SEM. \**p* < 0.05, \*\**p* < 0.01, \*\*\**p* < 0.001.

To provide a more detailed characterisation of the temporal extent of amnesia, we integrated both cohorts for a decade-by-decade analysis in a subset of participants who were able to provide personal events spanning one anterograde and six retrograde decades. Thirty-six participants were included in the analysis (N = 36): 18 with amnesia in the subset integrated cohort (3.0-Tesla cohort, n = 8; 7.0-Tesla cohort, n = 10) and their 18 matched controls from their respective cohorts. A mixed-model ANOVA 2 (group) by 2 (detail type) by 7 (time period: last year; 50–59, 40–49, 30–39, 20–29, 10–19, 0–9 years) again revealed significant main effects of group (*F*_(1,34)_ = 32.21, *p* < 0.001, η² = 0.49), time (*F*_(6,204)_ = 2.96, p = 0.009, η² = 0.08), and detail type (*F*_(1,34)_ = 440.12, *p* < 0.001, η² = 0.92), with group by time (*F*_(6,204)_ = 3.71, *p* = 0.002, η² = 0.10) and group by detail type (*F*_(1,34)_ = 42.07, *p* < 0.001, η² = 0.55) interactions but no three-way interaction (Figure 2C–D; Supplementary data 2). The earliest decade (0–9 years) showed no significant between group differences for episodic (*F*_(1,34)_ = 1.65, *p* = 0.21, η² = 0.04) or semantic details (*F*_(1,34)_ = 2.58, *p* = 0.12, η² = 0.07). Excluding this decade, a follow-up ANOVA confirmed a significant group by detail type interaction (*F*_(1,34)_ = 42.716, *p* < 0.001, η² = 0.557) but no group by time interaction (*F*_(5,170)_ = 0.89, *p* = 0.49, η² = 0.026). Planned comparisons (Bonferroni-corrected α = 0.025) revealed episodic details were significantly impaired (*F*_(1,34)_ = 45.07, *p* < 0.001, η² = 0.56), but semantic details remained intact (*F*_(1,34)_ = 4.18, *p* = 0.05, η² = 0.10). Together, these analyses demonstrate that focal hippocampal damage produces a selective loss of episodic autobiographical details extending over more than 50 years of life, with sparing confined to early childhood.

### Hippocampal CA2/3 volume predicts lifespan episodic retrieval

To examine structure-function relationships underlying the episodic amnesia, we tested whether hippocampal subfield volumes predicted autobiographical memory performance. Replicating previous findings ^11^, stepwise linear regression on the new 3.0-Tesla cohort revealed a significant model (*F*_(1,34)_ = 17.346, *p* < 0.001), with total CA2/3 volume selectively predicting internal details retrieved (*t* = 4.165, *p* < 0.001, R² = 0.338, β₁ = 0.581; see Figure 2E). Total CA1 volume (also reduced in volume in the new 3.0-Tesla cohort) volume was excluded from the model (*t* = -1.543, *p* = 0.132), even with reversed variable entry. The selective CA2/3 association was replicated with the extended Autobiographical Interview (7 life periods) in the integrated cohort (*F*_(1,34)_ = 9.031, *p* = 0.005). Total CA2/3 volume Z-score selectively predicted internal details (*t* = 3.005, *p* = 0.005, R² = 0.210, β₁=0.458; see Figure 2F), with CA1 volume excluded even with reversed variable entry (*t* = 1.074, *p* = 0.290). The large effect sizes in the new and subset integrated cohorts (R² = 0.338 and 0.210, respectively) demonstrate that CA2/3 volume contributes significantly when remembering personal episodic details from across the lifespan. Finally, a stepwise linear regression on the integrated cohort and controls (N = 64: 32 with amnesia (3.0-Tesla cohort, n = 18; 7.0-Tesla cohort, n = 14; 32 matched controls) revealed a significant model (*F*_(1,62)_ = 14.570, *p* < 0.001), with CA2/3 Z-score predicting total interval details (excluding 0-11 years) (*t* = 3.817, p<0.001 R² = 0.190, β₁ = 0.436), whereas the CA1 Z-score was excluded from the model (*t* = -0.435 *p* = 0.665).

Mediation analysis confirmed that the between-group difference in episodic memory performance was significantly mediated by CA2/3 volume in the new 3.0-Tesla cohort (N = 36; Sobel’s *Z* = 4.404, *p*<0.001, Cohen’s d = 0.734), demonstrating that CA2/3 damage directly contributes to the observed impairments. This mediation effect was replicated with the extended Autobiographical Interview in the subset integrated cohort (N = 36; Sobel’s *Z* = 4.447, *p* < 0.001, Cohen’s d = 0.786) and in the full integrated cohort (N = 64; Sobel’s *Z* = 4.215, *p* < 0.001, Cohen’s d = 1.054).

### Anterior-posterior CA3 gradients predict lifespan episodic retrieval

To investigate how functional specialisation along the hippocampal long axis manifests in episodic amnesia, we examined whether anterior-posterior structural variations predicted episodic retrieval performance (measured with the standard administration of the Autobiographical Interview) within the integrated cohort (N = 64): 32 with amnesia (3.0-Tesla cohort, n = 18; 7.0-Tesla cohort, n = 14) and 32 matched controls. For these analyses, we elected to use transformed anterior/posterior CA3 and CA1 volumes (*Z*-scored to account for individual variation in total hippocampal size and cohort; see Methods).

To characterise whether the broad CA3 contribution reflects anterior-posterior functional specialisation, we next examined whether anterior and posterior CA3 subdivisions demonstrated differential predictive power across memory time periods. Exploratory linear regression demonstrated that transformed anterior CA3 volume *Z*-scores predicted total internal detail scores for all memory time periods (11-18: *t* = 2.108, *p* = 0.039, R^2^ = 0.052, β₁= 0.259; 18-30: *t* = 2.082, *p* = 0.041, R^2^ = 0.050, β₁= 0.256; 30-55: *t* = 3.009, *p* = 0.004, R^2^ = 0.113, β₁= 0.357; last year: *t* = 3.465, *p*<0.001, R^2^ = 0.149, β₁= 0.403), except for the 0-11 time period (*t* = 0.385, *p* = 0.721, R^2^ = -0.014, β₁= 0.045). Sobel’s test demonstrated that there was a significant indirect effect of group membership mediating the relationship between transformed anterior CA3 *Z*-score and the cumulative internal (episodic) detail score (see Supplementary data 3). Transformed posterior CA3 volumes predicted total internal detail scores for the 18-30 (*t* = 2.488, *p* = 0.016, R^2^ = 0.076, β₁= 0.301), 30-55 (*t* = 3.260, *p* = 0.002, R^2^ = 0.146, β₁= 0.383), and last year time periods (*t* = 3.278, *p* = 0.002, R^2^ = 0.148, β₁= 0.384), but not 0-11 (*t* = 0.065, *p* = 0.949, R^2^ = -0.016, β₁ = 0.008) or 11- 18 (*t* = 1.776, *p* = 0.081, R^2^ = 0.033, β₁= 0.220) time periods. Sobel’s test once again demonstrated a significant indirect effect of group membership mediating the relationship between transformed posterior CA3 *Z*-score and the cumulative internal (episodic) detail score (see Supplementary data 3).

Anterior CA1 *Z*-scores were found to predict internal (episodic) detail score for the last year interval only (*t* = 2.325, *p* = 0.023, R^2^ = 0.080, β₁ = 0.283; see Supplementary data 3 for full results), with Sobel’s test demonstrating an indirect effect of group membership mediating this relationship. Posterior CA1 *Z*-scores were similarly found to predict internal (episodic) detail score for the last year interval (*t* = 2.799, *p* = 0.007, R^2^ = 0.112, β₁ = 0.335), with Sobel’s test demonstrating an indirect effect of group membership mediating the relationship between transformed posterior CA1 *Z*-score and the internal (episodic) detail score.

Together, the findings demonstrate a predictive role for CA3 in episodic retrieval and narrower, event-specific contribution of CA1, whereas anterior and posterior CA1 both specialised only for recent memories. The mediation through group indicates that these effects are functionally relevant to episodic amnesia rather than incidental volumetric variations.

### Hippocampal damage leads to content-specific vulnerability of lifespan episodic retrieval

To determine whether hippocampal damage selectively affects specific components of autobiographical memory content, we compared the integrated cohort with controls (N = 64) across five memory context-bound component types, characterised by the Autobiographical Interview (event details, temporal, place, perceptual, and emotion/thought). A mixed-model ANOVA (see Supplementary data 4 for full results and df corrections) revealed a significant main effect of group (*F*_(1,62)_ = 16.059, *p*<0.001, η² = 0.201) and significant interactions between group, memory detail type, and memory component (*F*_(2.33,144.33)_ = 12.574, p<0.001, η² = 0.164) and between group, memory detail type, memory component type, and time (*F*_(7.299,467.159)_ = 2.226, *p* = 0.029, η^2^ = 0.034). Post hoc one-way ANOVAs (α = 0.005) revealed no significant between-group differences for the earliest period (0–11 years) across internal event details (*F*_(1,62)_ = 1.043, *p* = 0.311, η^2^ = 0.016), temporal (*F*_(1,62)_ = 1.778, *p* = 0.187, η^2^ = 0.027), place (*F*_(1,62)_ = 0.041, *p* = 0.840, η^2^ = 0.001), perceptual (*F*_(1,62)_ = 0.036, *p* = 0.849, η^2^ = 0.001), and emotion/thought details (*F*_(1,62)_ = 3.502, *p* = 0.066, η^2^ < 0.052). This pattern extended to external event details (F statistics < 2.926, p > 0.092, η²s < 0.044; see Supplementary data 4).

A follow-up mixed-model 2 (group) by 2 (memory detail type: internal, external) by 4 (time period: last year, 30–55, 18–30, 11–18 years) by 5 (component type: event, time, place, perceptual, emotion/thought) ANOVA excluding the earliest period (0–11 years) revealed significant main effects of group (*F*_(1,62)_ = 20.822, *p* < 0.001, η^2^ = 0.245), memory detail type (*F*_(1,62)_ = 265.683 *p* < 0.001, η^2^ = 0.806), time period (*F*_(3,186)_ = 2.972, *p* = 0.033, η^2^ = 0.044), and component type (*F*_(1,62)_ = 408.803, *p* < 0.001, η^2^ = 0.865), with a significant four-way interaction (*F*_(12,768)_ = 2.150, *p* = 0.012, η² = 0.033; Supplementary data 4).

To characterise this four-way interaction suggesting temporal detail modulation, we conducted *post hoc* two-way ANOVAs examining group by time interactions for each internal detail component type. Only temporal details exhibited a significant group by time interaction (*F*_(3,192)_ = 3.907, *p* = 0.010, η² = 0.058; see Figure 3B and Supplementary data 4). Follow-up one-way ANOVAs across the four time periods for the time details (α = 0.0125) revealed that hippocampal damage produced the largest effect for recent memory (last year: *F*_(1,62)_ = 26.300, *p*<0.001, η² = 0.291). However, remote memory showed a non-monotonic pattern: temporal details from ages 11–18 showed greater hippocampal dependency (*F*_(1,62)_ = 5.923, *p* = 0.018, η² = 0.085) than from ages 18–30 (*F*_(1,62)_ = 0.660, *p* = 0.419, η² = 0.010) or 30–55 (*F*_(1,62)_ = 4.672, *p* = 0.034, η² = 0.068). This non-monotonic pattern indicates that hippocampal contributions to temporal detail retrieval vary across the lifespan, with maximal vulnerability for recent memories (Figure 3B), whereas time-invariant impairments were obtained across all of the other episodic memory components.

**Figure 3.**
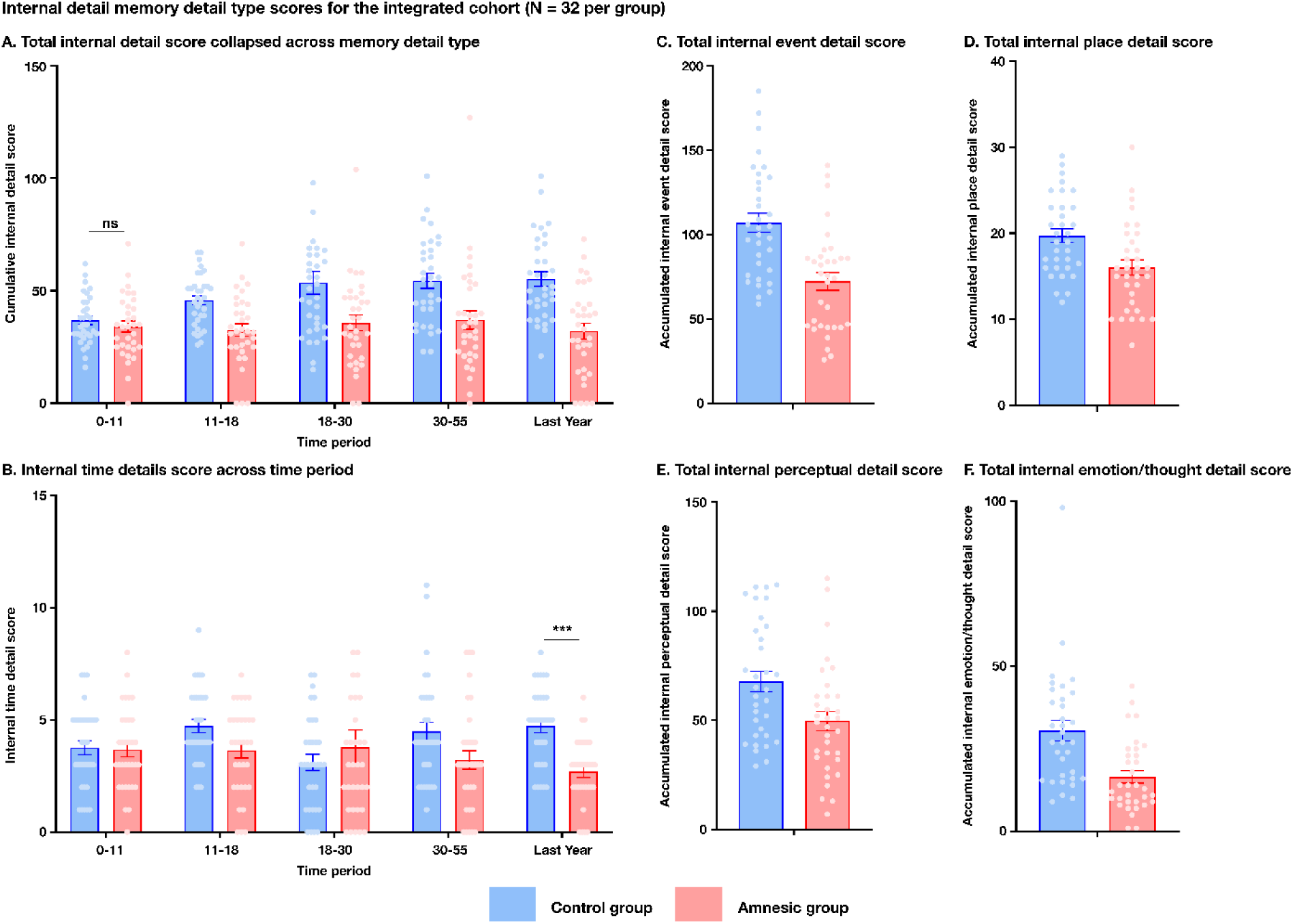
Content-specific episodic detail impairments across the lifespan in hippocampal amnesia (n = 32 per group). (A) Internal (episodic) detail scores across five life periods (0–11, 11–18, 18–30, 30–55 years, and last year) in the integrated cohort and controls (n = 32 per group; N = 64), showing significant group differences for all life periods except for the earliest childhood period (0–11 years) across all content categories (*F’*s<3.502, *p*’s > 0.066). (B) Temporal details (temporal information within remembered events) across life periods. A significant group by time period interaction (*F*_(3,186)_ = 3.907, *p* = 0.010, η² = 0.058) revealed a non-monotonic pattern that contrasts with other memory content categories, with impairment greatest for recent memories (last year: *F*_(1,62)_ = 26.300, *p* < 0.001, η² = 0.291). (C) Event details (specific happenings within remembered events) across life periods. A significant main effect of group (*F*_(1,62)_ = 23.413, *p* < 0.001, η² = 0.268) but no group by time period interaction (*F*_(3,186)_ = 1.381, *p* = 0.250, η² = 0.021) demonstrated time-invariant impairment across post-early-childhood periods. (D) Place details (spatial location information) across life periods. A significant main effect of group (*F*_(1,62)_ = 17.138, *p* < 0.001, η² = 0.211) but no group by time period interaction (F_(3,186)_ = 0.945, p = 0.420, η² = 0.015) demonstrated time-invariant impairment. (E) Perceptual details (sensory information) across life periods. A significant main effect of group (*F*_(1,64)_ = 10.301, *p* = 0.002, η² = 0.139) but no group by time period interaction (*F*_(3,192)_ = 0.187, *p* = 0.667, η² = 0.003) demonstrated time-invariant impairment. (F) Emotions/thoughts (emotional and cognitive content) across life periods. A significant main effect of group (*F*_(1,62)_ = 14.286, *p* < 0.001, η² = 0.182) but no group by time period interaction (*F*_(3,186)_ = 0.705, *p* = 0.522, η² = 0.011) demonstrated time-invariant impairment. Panels C–F demonstrate that hippocampal damage produces time-invariant deficits across event, place, perceptual, and emotional/cognitive content, whereas temporal details (Panel B) show non-monotonic vulnerability. Each dot represents a single participant (red, group with amnesia; blue, control group). Full statistical analyses are reported in the main text and Supplementary Tables. Error bars represent SEM. *p < 0.05, **p < 0.01, ***p < 0.001. ns = not significant.

### CA3 and CA1 mediate distinct episodic detail components

In the integrated cohort with amnesia and controls (N = 64), linear regression analyses revealed that total transformed CA3 volume *Z*-scores significantly predicted each context-bound component comprising the internal (episodic) detail score (excluding the 0-11 period internal detail subcategories): event (*t* = 4.256, *p*<0.001, R^2^ = 0.226, β₁= 0.476), place (*t* = 2.565, *p* = 0.013, R^2^ = 0.096, β₁= 0.310), temporal (*t* = 2.777, *p* = 0.007, R^2^ = 0.096, β₁= 0.333), perceptual (*t* = 2.481, *p* = 0.016, R^2^ = 0.090, β₁= 0.300), emotion/thought details (*t* = 3.346, *p* < 0.001, R^2^ = 0.162, β₁= 0.403) and total internal (episodic) details (*t* = 4.242, *p* < 0.001, R^2^ = 0.212, β₁ = 0.474). Mediation analyses demonstrated significant indirect effects of group membership on the relationship between transformed CA3 *Z*-score and the dependent variable (see Supplementary data 5 for full results). Similar results were obtained modelling total internal (episodic) memory detail type score with CA2/3 volume *Z*-scores and the memory detail types (including mediation analyses; see Supplementary data 5).

In contrast, total CA1 volume *Z*-score predicted only total internal detail (*t* = 2.084, *p* = 0.041, R^2^ = 0.065, β₁= 0.256) and event detail performance (*t* = 2.210, *p* = 0.031, R^2^ = 0.73, β₁= 0.270), with non-significant associations for place, time, perceptual, and emotion/thought categories (see Supplementary data 5). Mediation analyses revealed significant indirect effects of group membership on the relationship between transformed CA1 *Z*-score and both total internal (*Z* = 3.402, *p* = 0.002, Cohen’s d = 0.425) and event detail scores (*Z* = 3.395, *p* = 0.001, Cohen’s d = 0.424).

### Local narrative coherence but not global narrative coherence is impaired by hippocampal damage

To assess whether hippocampal damage affects the organisational structure of autobiographical memory narratives, we analysed local and global narrative coherence in the integrated cohort of individuals with amnesia and the matched controls (N = 64). A mixed-model 2 (group: amnesic, control) by 5 (time period: last year, 30–55, 18–30, 11–18, 0–11 years) ANOVA on local narrative coherence scores revealed a significant main effect of group (*F*_(1,62)_ = 4.251, *p* = 0.043, η^2^ = 0.064; Figure 4A) but not time period (*F*_(3.51,217.38)_ = 0.835, *p* = 0.504, η^2^ = 0.013). There was no significant two-way interaction between group and time period (*F*_(3.51,217.38)_ = 1.817, *p* = 0.141, η^2^ = 0.028) indicating a time-invariant reduction in local narrative coherence following hippocampal damage compared to the controls. This pattern of results was repeated when the most remote decade was excluded (see Supplementary data 6 for full results and df correction). Crucially, the corresponding analysis for global narrative coherence showed no significant group effect (*F*_(1,62)_ = 0.790, *p* = 0.378, η^2^ = 0.013), indicating preserved global coherence despite hippocampal damage (Figure 4B; Supplementary data 6).

**Figure 4.**
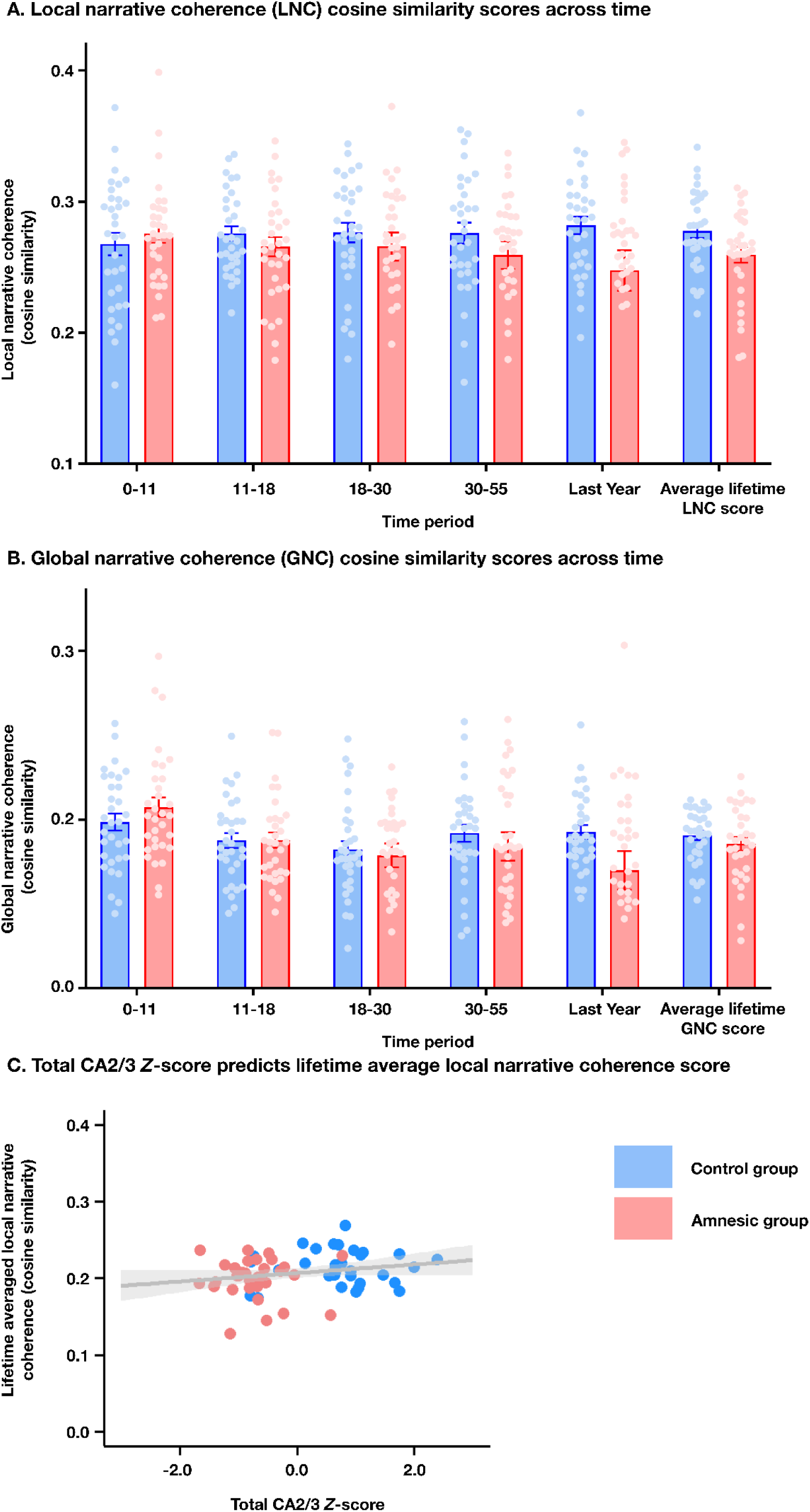
Hippocampal CA2/3 damage impairs local narrative coherence but preserves global narrative coherence across the lifespan. (A) Local narrative coherence (mean cosine similarity between adjacent sentence embeddings) across five life periods (0–11, 11–18, 18–30, 30–55 years, and last year) in the integrated cohort (n = 32 individuals with amnesia; n= 32 controls; N = 64). A significant main effect of group (*F*_(1,34)_ = 6.327, *p* = 0.017, η^2^ = 0.157) but no group by time period interaction (*F*_(4,136)_ = 0.352, *p* = 0.843, η^2^ = 0.010) demonstrated reduced local coherence in the group with amnesia compared to controls. (B) Global narrative coherence (mean cosine similarity between each sentence and the narrative centroid) across the same life periods. No significant main effect of group was observed, indicating that global narrative coherence remained intact despite hippocampal damage. (C) Total CA2/3 volume Z-score (independent variable) predicts lifetime-averaged local narrative coherence (dependent variable, excluding 0–11 years) in the integrated cohort. Stepwise linear regression revealed a significant model (*F*_(1,64)_ = 4.203, *p* = 0.045), and a selective association between CA2/3 volume and local narrative coherence (*t* = 2.050, *p* = 0.045, R² = 0.063, β₁ = 0.252), whereas CA1 volume did not contribute significantly to the model (*t* = -0.806, *p* = 0.423). Each dot represents a single participant (red, group with amnesia; blue, control group), with the best-fitting linear regression line shown. The dissociation between impaired local narrative coherence (panel A) and preserved global narrative coherence (panel B) demonstrates that CA2/3 damage selectively disrupts fine-grained sequential binding of adjacent episodic elements while sparing broader narrative organisation. Full statistical analyses are reported in the main text and Supplementary Tables. Error bars represent SEM. *p < 0.05, **p < 0.01, ***p < 0.001. ns = not significant.

Next, to determine whether reduced local coherence reflected random or variable narrative production rather than selective impairment in sequential semantic binding, we examined two complementary metrics derived from local narrative coherence: (1) variance (consistency of semantic binding) and (2) entropy (information-theoretic predictability of sequential transitions via Shannon entropy). Variance and entropy showed no significant between-group differences (variance: *F*_(1,62)_ = 0.009, *p* = 0.925, η^2^ = 0.000; entropy: *F*_(1,62)_ = 0.234, *p* = 0.630, η^2^ = 0.004), indicating preserved consistency and predictability of narrative transitions. Hence, reduced local narrative coherence together with intact global coherence reflects a selective impairment in sequential binding within remembered events and preserved broader organisation of narratives.

### Local narrative coherence mediates CA2/3 damage effects on episodic detail retrieval

To determine how this disruption in local narrative coherence relates to episodic memory performance across the lifespan, we next determined whether lifespan-averaged local narrative coherence scores predicted total episodic retrieval performance and components of episodic detail. In the integrated cohort and controls (N = 64), we found that lifespan-averaged local narrative coherence score predicted total (excluding the 0-11 time period) episodic (internal) detail performance (*t* = 5.059, *p*<0.001, R² = 0.292, β₁ = 0.541; see Figure 4C), whereas lifespan-averaged local narrative coherence did not predict total semantic (external) detail performance (*t* = 1.077, *p* = 0.289, R² = 0.033, β₁ = 0.182), thereby demonstrating specificity for hippocampal-dependent episodic processes. Mediation analysis confirmed that reduced local narrative coherence significantly mediated the relationship between CA2/3 damage and episodic detail loss (Sobel’s Z = 2.064, *p* = 0.039, Cohen’s d = 0.383), establishing local narrative coherence as a mechanistic pathway linking CA2/3 damage to episodic memory impairment. The selective vulnerability of local but not global narrative measures provides internal convergent validity and demonstrates that the observed group differences reflect dissociable aspects of narrative organisation that depend on hippocampal CA2/3 integrity rather than measurement artifact or individual differences in narrative style.

Local narrative coherence demonstrated component-specific predictive power for internal event (*t* = 3.611, *p*<0.001, R² = 0.174, β₁ = 0.417), place (*t* = 3.589, *p*<0.001, R² = 0.172, β₁ = 0.415), perceptual (*t* = 5.596, *p*<0.001, R² = 0.336, β₁ = 0.579), emotions/thoughts (*t* = 3.161, *p* = 0.002, R² = 0.139, β₁ = 0.373), and temporal details (*t* = 2.400, *p* = 0.019, R² = 0.085, β₁ = 0.292; see Figures 5A-F). Corresponding mediation analyses revealed significant indirect effects of group membership for internal event (Sobel’s *Z* = 2.261, *p* = 0.024, Cohen’s d = 0.283), place (Sobel’s *Z* = 2.160, *p* = 0.031, Cohen’s d = 0.270), perceptual (Sobel’s *Z* = 1.978, *p* = 0.048, Cohen’s d = 0.247), and emotional/cognitive (Sobel’s *Z* = 2.103, *p* = 0.035, Cohen’s d = 0.263), but not for temporal details (Sobel’s *Z* = 1.785, *p* = 0.074, Cohen’s d = 0.223).

**Figure 5.**
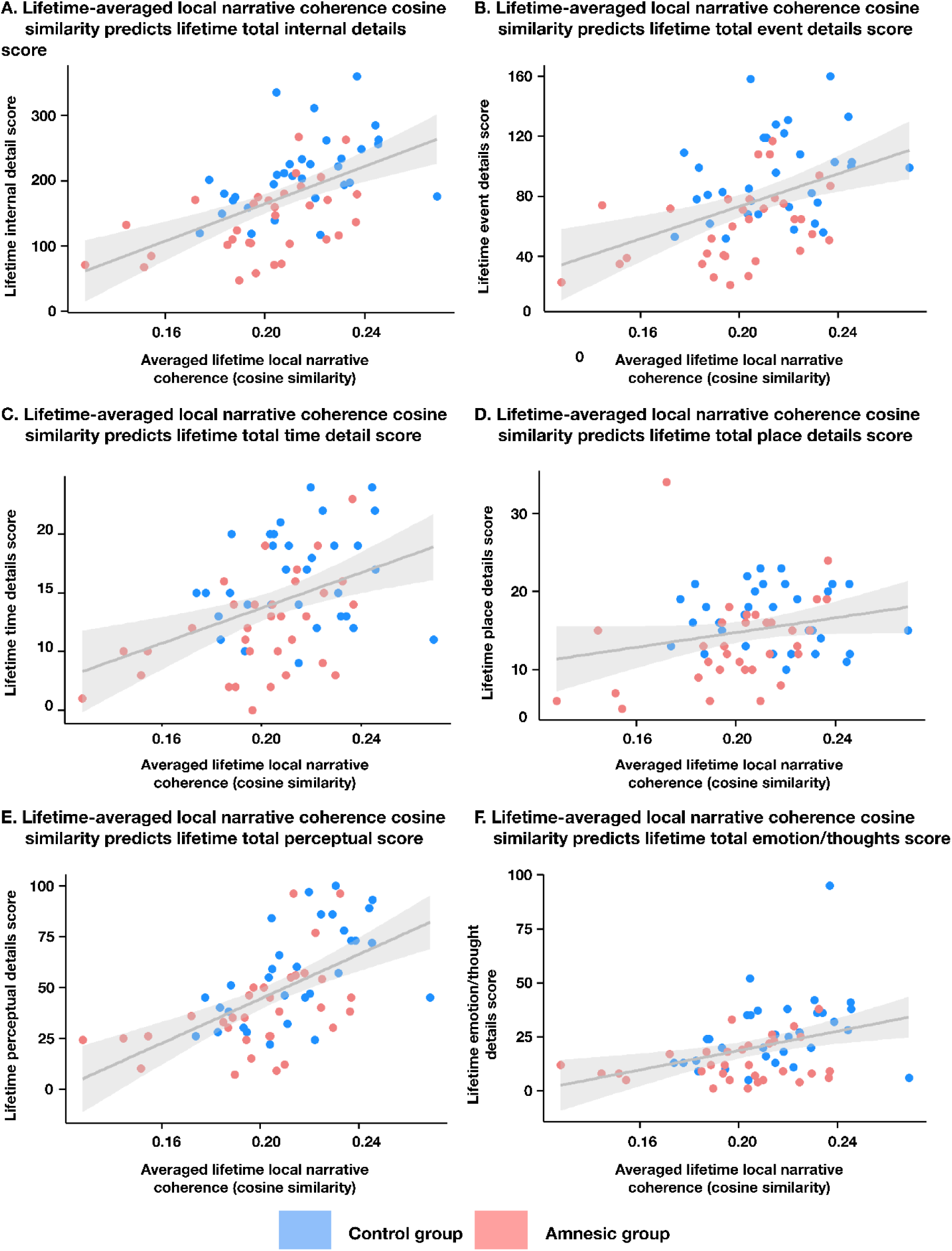
Local narrative coherence predicts episodic detail performance across context-bound categories of memory content. (A) Lifetime-averaged local narrative coherence (independent variable, excluding 0–11 years) predicts lifetime-averaged internal (episodic) detail scores (dependent variable, excluding 0–11 years) in the integrated cohort and controls (n = 32 individuals with amnesia; n= 32 controls; N = 64). Linear regression revealed a significant association (*t* = 5.059, *p*<0.001, R² = 0.292, β₁ = 0.541), whereas local narrative coherence did not predict external (semantic) details (*t* = 1.077, *p* = 0.289, R² = 0.033, β₁ = 0.182), demonstrating detail type specificity. (B–F) Local narrative coherence also predicted specific internal memory content categories: (B) event details (*t* = 3.611, *p*<0.001, R² = 0.174, β₁ = 0.417); (C) place details (*t* = 3.589, *p*<0.001, R² = 0.172, β₁ = 0.415); (D) temporal details (*t* = 2.400, *p* = 0.019, R² = 0.085, β₁ = 0.292); (E) perceptual details (*t* = 5.596, *p*<0.001, R² = 0.336, β₁ = 0.579); and (F) emotions/thoughts (*t* = 3.161, *p* = 0.002, R² = 0.139, β₁ = 0.373). Each panel shows the relationship between local narrative coherence and the respective context-bound memory content category. These findings demonstrate that impaired local narrative coherence within remembered events predicts episodic detail loss across all content categories, establishing a mechanistic link between narrative organisation and memory performance. Each dot represents a single participant (red, group with amnesia; blue, control group), with the best-fitting regression line shown. Full statistical analyses are reported in the main text and Supplementary data.

Finally, we examined whether local narrative coherence disruption in the integrated cohort (excluding the most remote period) could be predicted by CA2/3 volume or CA1 volume *Z-*scores using a stepwise linear regression. A significant model was found (*F*_(1,64)_ = 4.203, *p* = 0.045). Total CA2/3 volume *Z*-scores significantly predicted local narrative coherence (lifespan-averaged) scores (*t* = 2.050, *p* = 0.045, R² = 0.063, β₁ = 0.252; see Figure 4C), whereas total CA1 volume *Z*-scores were excluded from the model, even when the entry of variables was reversed (*t* = -0.806, *p* = 0.423).

## Discussion

Fundamental questions remain about how human hippocampal subfield specialisation contributes to autobiographical memory across lifespan consolidation, the retrieval of context-bound components of episodic memory, and the narrative organisation of remembered events. We studied 32 individuals with amnesia, secondary to focal, bilateral hippocampal damage, and obtained causal evidence that links subfield anatomy to these three aspects of episodic retrieval. Across six decades of remembered events, we observed a time-invariant loss of episodic detail except early childhood (0–11 years) and the sparing of semantic details. CA2/3 volume predicted episodic retrieval and all context-bound components of episodic memory (internal detail, event specifics, place, temporal information, perceptual content and emotion/thought), whereas CA1 volume predicted only recent memories and event details. Computational linguistic measures of how episodic details were linked within remembered events, assessed via sentence-level SBERT, demonstrated that local narrative coherence (mean cosine similarity between adjacent sentence embeddings) mediated episodic detail loss across multiple content domains in the cohort with hippocampal amnesia, with CA2/3 damage leading to impaired local narrative coherence, whereas global narrative coherence was preserved.

The concordant profile of time-invariant episodic impairment in the hippocampal CA2/3 cohort across behavioural and computational-linguistic measures supports hybrid models of episodic memory consolidation, wherein neocortical networks represent schematic traces whilst the hippocampus remains essential for context-rich recollection across the lifespan ^3,4,68,69^. Recent non-human animal evidence has also corroborated these models, showing that contextual reminders reinstate hippocampal engrams and override cortical generalisation ^70^, and that persistent hippocampal–cortical dialogue is required to sustain episodic detail ^28^. The resilience of early-life memories observed in the current study likely reflects heightened developmental neurogenesis and the establishment of neocortical schemas before full maturation of hippocampal–cortical networks ^28,71^. Critically, CA3-dependent neurogenesis is necessary for post-childhood memories but not for the earliest memories ^72^, which suggests that early childhood memories persist through schema-based, cortex-dependent mechanisms distinct from those underlying adult hippocampal-dependent episodic retrieval ^71,73,74^.

The time-invariant loss in the cohort with hippocampal damage across most context-bound components of episodic memory (i.e., event specifics, place, perceptual features, and emotional/cognitive content), with the notable differential vulnerability in context-bound temporal details, underscores the role of the hippocampus in binding multimodal contextual elements into coherent episodic memories. Hierarchical binding theories propose that the hippocampus integrates medial prefrontal schemas, posteriomedial contextual codes, and anterotemporal object/person representations to support vivid first-person recall ^2,30,50,69,71,75–77^, which recent computational models suggest involves hippocampal replay training generative models to recreate sensory experiences from latent variable representations ^4^. The differential vulnerability of temporal details relative to the control group, which was maximal in recent memories, minimal in young adulthood (18-30 years), and intermediate in early adolescence (11-18 years) and mid-adulthood (30-55 years), likely reflects changes in the demands on hippocampal-mediated contextual binding across retention intervals. Recent memories require active temporal sequencing, whereas older memories benefit from cortical reorganisation of temporal structure into schema-based representations. These findings support frameworks that posit selective and graded hippocampal contributions to specific memory components ^78,79^, challenging classical models of uniform episodic impairment following hippocampal injury ^69^.

Computational linguistic analyses revealed the predicted dissociation: hippocampal damage impaired local narrative coherence within remembered personal events, whereas global narrative coherence remained intact. Notably, the time-invariant reduction in local narrative coherence across post-early-childhood memories corresponds with the episodic detail loss pattern that spared only early childhood. Mediation analyses demonstrated that reduced local narrative coherence significantly mediated hippocampal CA2/3 damage effects on episodic detail retrieval, specifically for event details, place, perceptual, and emotional/cognitive content but not temporal details, thereby establishing local narrative coherence as a critical pathway through which CA2/3 damage compromises episodic richness. Crucially, reduced mean local narrative coherence co-occurred with intact variance and entropy, which indicates preserved distributional properties and predictability of sequential transitions within remembered events. This dissociation demonstrates that hippocampal CA2/3 mediates the strength of sequential bindings rather than the overall organisational scaffold of episodic narratives.

The selective impact on binding strength is consistent with computational models that propose CA3 mediates associative strength between sequential elements rather than the overall scaffold of episodic representations ^2,33,47,52,80^. Consecutive narrative sentences contain overlapping contextual features (temporal proximity, character continuity, spatial setting) that require disambiguation for local coherence; the recurrent architecture of CA3 contributes to this operation through pattern separation and completion operations ^81,82^. By contrast, CA1 supports gist-based memories and integration across broader narrative contexts through hippocampal-neocortical interactions ^50,51^, paralleling the distinction between within-event sequential binding and cross-event associative integration ^83^. Consistent with this framework, theta power and hippocampal–parahippocampal connectivity scale with semantic distances between consecutively remembered items ^84^. However, focal hippocampal damage produces distal network effects on medial temporal and frontal connectivity ^11,30^, such that the extent to which impaired local narrative coherence reflects primary CA2/3 circuit dysfunction versus downstream hippocampal–cortical disruption remains to be determined.

Prior investigations of narrative structure in amnesia have employed holistic discourse ratings ^85,86^ or assessed whether details from externally-presented narratives were remembered in sequence ^87^. Related work has examined hippocampal integration across event boundaries ^48,83,88,89^ and applied natural language processing to autobiographical narratives ^26,90^, though these approaches focused on event segmentation, schematic flow, or neural reactivation rather than quantifying subfield-level coherence mechanisms. Computational coherence frameworks have been validated in aging and dementia populations ^44–46^, but have not been applied to focal hippocampal subfield damage. By applying SBERT-based metrics ^42,43^, the present study demonstrates that focal hippocampal damage selectively disrupts self-generated sequential binding whilst sparing broader narrative coherence. Future work should establish whether this dissociation generalises to non-autobiographical episodic narratives ^89,91^ and other populations with hippocampal pathology, determining whether local narrative coherence captures a domain-general principle of hippocampal contributions to episodic memory organisation. In parallel, it will be important to relate these coherence mechanisms to boundary- and context-sensitive structuring of recall and continuity observed in recent behavioural paradigms ^92,93^.

Volumetric analyses revealed distinct functional profiles along the hippocampal long axis. CA2/3 volume predicted episodic detail retrieval across all post-childhood life periods and mediated group differences across multiple detail categories. Anterior CA3 predicted detail retrieval across the broadest temporal range (11–55+ years), whereas posterior CA3 showed selective engagement from young adulthood onwards (18–55+ years), consistent with computational models linking anterior hippocampus to associative binding ^32,53,94^. In contrast, total and anterior–posterior subdivisional CA1 volumes predicted only recent memories (last year) and event specifics, reflecting distinct computational demands: CA1 mediates discrimination of recent, interfering memories through rapid temporal resolution and pattern separation, whereas CA3 supports remote memories through autoassociative reinstatement of item–context bindings ^2,35,36^. Mechanistically, posterior CA1 orthogonalises perceptually similar recent environments via visual–spatial circuits, whereas anterior CA1 resolves conceptual overlap between new episodes and existing schemas through anterior–temporal and prefrontal networks ^71,95^. As memories age, competing traces fade or become semanticised, reducing CA1-mediated discrimination demands and explaining the recent-only CA1 effect without invoking consolidation-driven memory relocation.

The current findings challenge the temporal gradient predicted by systems consolidation theory, yet non-human animal models of hippocampal lesions continue to yield graded retrograde amnesia ^96–99^. In rodent contextual fear conditioning, CA3-mediated pattern completion declines as memories consolidate to gist-like representations, no longer requiring CA3 recurrent networks ^33,34,100^, whereas CA1 can support recent and remote memories ^101,102^. However, high-fidelity tasks such as spatial navigation ^103^ or autobiographical recall ^11^ re-engage CA3-dependent auto-association because perceptual, spatial, and emotional details require binding. Within the contextual binding framework ^2^, CA1 serves dual roles: discriminating recent events under temporal interference ^35,36^ and maintaining gist-level retrieval across all timescales ^102^. However, detailed retrieval, requiring perceptual, spatial, and emotional binding, shows declining CA1 engagement as memories age. Remote gist-level memories show reduced CA1 Arc expression ^103^, and CA1 damage in humans produces a temporal gradient with maximal deficits for recent detailed memories ^104^, whereas CA3 maintains persistent engagement when episodic detail is required, regardless of memory age. Thus, the presence or absence of temporal gradients appears to depend on whether retrieval demands access gist-level representations or high-fidelity episodic details.

Task-species differences also help to explain divergent findings: non-human paradigms typically employ paired-associate or avoidance learning ^105,106^, which consolidate to gist representations over time. Here we assessed self-relevant, multisensory autobiographical memories that recruit persistent binding mechanisms due to their complexity and personal salience. Furthermore, the human hippocampus integrates extensively with heteromodal networks including the default mode network (DMN), whereas non-human primates exhibit incomplete DMN integration ^107,108^. Following hippocampal damage, humans rely more heavily on distributed neocortical reorganisation than rodents ^71,109^, and hippocampal neurogenesis produces rapid memory consolidation in mice ^97^, indicating that temporal gradients arise from active circuit-level processes rather than passive decay. These species-specific differences in network architecture and consolidation dynamics may explain why human autobiographical memory shows time-invariant CA3 dependence, whereas animal conditioning paradigms yield temporally graded amnesia.

Several considerations contextualise these findings. First, the cross-sectional design does not track individual memory trajectories over time; temporal patterns represent between-participant comparisons across life periods rather than longitudinal changes within the same individuals. Second, the computational coherence metrics assume that narrative structure reflects underlying memory organisation. The variance and entropy controls help to separate retrieval deficits from impairments in narrative production, but this distinction cannot be fully disentangled. Third, autobiographical episodic memories are characterised by personal salience, multimodal binding, and self-referential processing, and thus may not generalise to simpler laboratory-based episodic tasks that do not share these features. Finally, while the single aetiology (LGI1-LE) provides mechanistic clarity by isolating focal CA2/3 damage, generalisability to other amnesia aetiologies (e.g., Alzheimer’s disease, traumatic brain injury) with differing lesion profiles or progressive pathology remains an empirical question.

In summary, hippocampal CA2/3 damage produces selective and durable vulnerabilities in episodic detail retrieval and narrative organisation that persist across lifespan memories, challenging classical systems consolidation accounts and supporting frameworks that emphasise parallel, independent hippocampal–cortical contributions ^110^. The time-invariant impairment in local narrative coherence alongside preserved global coherence reveals that episodic memory reconstruction operates through distinct computational processes at different organisational scales, with CA2/3 specifically contributing to fine-grained contextual binding between adjacent narrative elements. The mediation of episodic detail loss through impaired local coherence, combined with non-monotonic vulnerability of temporal details, indicates that CA2/3 contributions are both content-specific and mechanistically dissociable from broader integration supported by distributed hippocampal–cortical networks. Although episodic and semantic memory share neural substrates ^111^, both rely on distinct hippocampal-dependent computational processes. Local narrative coherence, selectively predicting episodic (but not semantic) detail retrieval, exemplifies one such process, demonstrating mechanistic dissociation despite substrate overlap. CA1 demonstrates preferential involvement in recent (< 1 year) rather than remote autobiographical memory, aligning with evidence that recently acquired memories are decoded within CA1 whilst remote memories are preferentially represented in posterior CA3 and dentate gyrus ^112,113^. Together, these results support a transition towards mechanistic, subfield-informed frameworks for understanding human episodic memory and memory dysfunction in hippocampal pathology.

## Methods

### Participants

All participants had chronic amnesia induced by a single-aetiology, leucine-rich glioma-inactivated-1 limbic encephalitis (LGI1-LE)^114^. LGI1-LE is typically a monophasic illness associated with static postmorbid cognition^11,29,115^. Accordingly, none of the participants with amnesia had clinical evidence of a relapse at any time surrounding or subsequent to the experimental period. The new cohort, characterised using 3.0-Tesla MRI, comprised 18 participants (mean age and SEM: 58.1±2.33 years; 13 male, 5 female) who underwent comprehensive neuropsychological assessment and the standard administration of the AI^30^. Eighteen healthy age- and education-matched controls were recruited (58.1±2.33 years; 13 male, 5 female) to the control group. A subset of 14 participants (mean age and SEM: 66.7±2.21 years; 11 male, 3 female) from our original cohort, characterised using 7.0-Tesla MRI, was selected based on availability of memories spanning at least six decades (0–9, 10–19, 20–29, 30–39, 40–49, 50–59, and last year), alongside 14 healthy age- and education-matched controls (mean age and SEM: 63.8±2.52 years; 12 male, 2 female) ^11^. All control participants had no history of cognitive, psychiatric, or neurological illness.

For the extended administration of the Autobiographical Interview, 32 participants with amnesia and 32 matched controls from both cohorts (eight participants from the 3.0-Tesla cohort and their matched controls; 10 participants from the 7.0-Tesla cohort and their matched controls) provided memories spanning at least six decades (group with amnesia mean age and SEM: 67.9±1.42 years; control group mean age: 66.1±1.38 years). For these participants, analyses of the standard administration of the AI used the memory with the highest internal detail count from the life periods 30–39, 40–49, or 50–59 to represent the 30–55 epoch. We then combined participants from both cohorts (32 per group; amnesia: mean age ± SEM = 61.1±1.83 years; controls: mean age ± SEM = 59.9±1.80 years) to ensure sufficient power for assessing the internal (episodic) and external (semantic) detail categories assessed in the standard administration of the Autobiographical Interview (see below).

All participants were fluent, native English speakers. All participants were fluent native English speakers. All participants with hippocampal amnesia were otherwise healthy (self-reported), with no evidence of secondary gain or active psychopathology. Informed written consent was obtained from all participants for all procedures and for consent to publish, in accordance with the terms of approval granted by a national research ethics committee (22/NW/0088 and 04/Q0406/147) and the principles expressed in the Declaration of Helsinki.

### Hippocampal segmentation

Detailed descriptions of the 3.0- and 7.0-Tesla MRI parameters for structural acquisition, hippocampal subfield segmentation, and quality assurance (e.g., Dice similarity coefficients) are reported in our previous reports ^11,29,30^. Hippocampal subfield volumes were computed using two complementary approaches to enable harmonisation across the distinct 3.0- and 7.0-Tesla scanning and segmentation protocols. This cross-cohort harmonisation strategy addresses well-established challenges in multi-site neuroimaging studies where scanner differences can introduce systematic biases that confound between-group comparisons ^65,66^.

First, for the original 7.0-Tesla cohort ^11^, a composite CA2/3 volume was calculated as the sum of total CA3 and CA2 volumes, consistent with established anatomical models in which CA2 and CA3 operate as a functional subregion ^63,64^. Population-level Z-scores for total CA2/3 volume (left and right hippocampal subfields pooled) were then computed across both the 3.0-Tesla and 7.0-Tesla cohorts, which enabled direct comparison of datasets despite different acquisition protocols. Z-scores were calculated by subtracting the population mean (including all participants with hippocampal amnesia and control participants from both cohorts: 3.0-Tesla n = 36; 7.0-Tesla n = 28) from each total CA2/3 volume, then dividing by the respective population standard deviation. This method generates standardised metrics that adjust for cohort-level variance but preserve individual differences ^116^.

The second approach addresses the technical limitation that CA3 and CA2 could not be reliably delineated within the 3.0-Tesla dataset, which is a recognised challenge given the superior contrast-to-noise ratio and spatial resolution of 7.0-Tesla imaging for hippocampal subfield visualisation ^67,117^. To address this technical limitation, empirically-derived proportional contributions of CA3 and CA2 to the composite CA2/3 volume were estimated from the high-resolution 7.0-Tesla cohort. This allowed group- and region-specific estimation of the relative CA3 contribution within composite volumes, thus leveraging anatomical precision from the 7.0-Tesla cohort to inform analysis of lower-resolution data. This transformation was performed separately for anterior and posterior hippocampal regions to accommodate known functional and structural heterogeneity along the longitudinal axis ^14,63^. The anterior-posterior boundary was defined as the last slice containing the uncal apex, following established anatomical landmarks ^63^.

For the anterior hippocampus in the 7.0-Tesla cohort, the empirically-derived CA3:CA2/3 ratio was 0.730 ± 0.014 (median: 0.732) for controls and 0.629 ± 0.014 (median: 0.633) for participants with hippocampal amnesia. In the posterior hippocampus, these ratios were 0.439 ± 0.015 (median: 0.427) for controls and 0.397 ± 0.014 (median: 0.401) for the group with amnesia. These ratios were then used to estimate the CA3 component, contributing to each composite CA2/3 volume in the 3.0-Tesla cohort. Separate ratios were applied for participants in the control group and group with amnesia to reflect group-specific differences, and region-specific transformation ratios were used for anterior versus posterior hippocampal distributions. The resulting CA3 volume estimates were then Z-scored using intra-cohort control means and standard deviations ^67^.

It should be noted that this estimation process assumes that group-level proportionality observed at 7.0-Tesla holds for the broader cohort, recognising the limitation that individual subfield variability cannot be perfectly resolved in lower-resolution imaging. However, this harmonisation approach enables standardised, cross-cohort comparisons while maximally exploiting the anatomical precision available in the reference dataset. These harmonised anterior and posterior CA3 volume Z-scores, alongside unaltered CA1 volume Z-scores from both cohorts, were used as independent variables in subsequent analyses, thus enabling cross-cohort structure-function investigations despite technical constraints of lower-field imaging.

### Neuropsychological assessment

Neuropsychological assessment was conducted in all participants using the following standardised subtests: *Verbal and visual intelligence:* Wechsler Abbreviated Scale of Intelligence (WASI) – Similarities and Matrix Reasoning^118^; *Premorbid intelligence:* Test of Premorbid Functioning – UK Version^119^; *Verbal memory:* Logical memory I and II from Wechsler Memory Scale–III [(WMS-III)^120^; *Visual memory:* Rey complex figure – Immediate-Recall and Delayed-Recall^121^; *Recognition memory*: Recognition Memory Test – Words and Faces^122^; *Sustained Attention*: Test of Everyday Attention – Map Search^123^; *Language*: Graded Naming Test ^124^; Letter Fluency and Category Fluency from the Verbal Fluency from Delis-Kaplan Executive Function System (D-KEFS)^125^; *Executive function:* Category Switching from the Verbal Fluency Test, Number-Letter Switching from the Trail Making Test from Delis-Kaplan Executive Function System (D-KEFS)^125^; and, WMS-III – Digit Span^120^; *Attentional Switching / Cognitive Flexibility:* Visual Elevator from Test of Everyday Attention^123^; *Visuomotor skills:* Visual Scanning, Number Sequencing, Letter Sequencing, and Motor Speed from the Trail Making Test from Delis-Kaplan Executive Function System (D-KEFS)^125^; and *Visuoconstruction skills*: Rey complex figure – Copy^121^. Scores on the standardised neuropsychological tests were first transformed into age-corrected standard values, where available, and then converted to Z-scores. One participant in each cohort with hippocampal amnesia was unable to complete the premorbid intelligence test (TOPF and NART) due to severe dyslexia.

### Neuropsychology statistical analyses

Data from the neuropsychological assessments were analysed using two-sample (two-tailed) t-tests to determine whether performance in the group with hippocampal amnesia was significantly different from normative data (mean = 0, SD = 0.3; representing age-corrected Z-scores).

### Autobiographical interview

The monophasic nature of autoimmune encephalitis indicates that hippocampal function was intact in all individuals with hippocampal amnesia during the encoding of memories formed prior to disease onset. Consequently, the observed retrograde memory impairments can be attributed to the disruption of hippocampal-mediated retrieval mechanisms rather than difficulties in anterograde encoding or consolidation ^11,29^. Episodic (denoted as internal details) and context-independent semantic (external details) memory were assessed using the Autobiographical Interview (AI)^31^, a semi-structured interview instrument that enables objective, parametric, text-based analysis of autobiographical recall. All participants completed the standard AI, which probes memories from five life periods: 0–11, 11–18, 18–30, 30–55 (retrograde periods), and last year (anterograde period; postmorbid for participants with amnesia). For decade-by-decade analyses, the AI was adapted for those able to provide one memory per decade, encompassing six retrograde decades plus one anterograde time point (last year), for a total of seven time periods: 0–9, 10–19, 20–29, 30–39, 40–49, 50–59 (retrograde), and last year (anterograde).

### Scoring and reliability

All verbal responses during the AI were recorded in digital audio format and then transcribed for offline scoring. Transcripts were de-identified such that any identifying personal details or references to group membership were removed. Responses were scored according to the standardised method outlined in the AI Scoring Manual ^31^. Each remembered event was segmented into informational units (details) comprising occurrences, observations, or thoughts expressed as a grammatical clause. These details were assigned to five qualitative response categories of information: event details, time, place, perceptual details, and emotions and thoughts. Details that related directly to a unique event, with a specific time and place or associated with episodic re-experiencing (e.g., thoughts or emotions), were classified as internal (episodic) details. Information that did not relate to the specific event was assigned to external details and sub-categorised into semantic (factual information or extended events), repetitions (previously stated details with no new elaboration), and other (e.g., metacognitive statements, editorialising, inferences). In keeping with convention ^126^, the internal (episodic) and external (semantic) point scores were accumulated across all three levels of cueing that formed the basis of the standard administration of the AI^31^. Composite internal and external detail scores were computed for every life epoch and participant by summating the scores across the five response categories for the standard AI administration (new 3.0-Tesla cohort) and extended AI administration (original 7.0-Tesla cohort). For the combined cohort analyses, internal and external scores were analysed as individual response categories (five categories by two detail types = 10 response category scores per epoch).

For the new 3.0-Tesla cohort, one rater (TM) scored 100% of the transcripts, with a second rater (JZ) independently scoring 50% for reliability purposes. For the original 7.0-Tesla cohort, two raters (including TM) scored 100% of the acquired episodes. This approach aligns with prior studies ^31,58^. Intra-class correlation coefficients (ICCs) using a two-way mixed-model design assessed absolute agreement between raters. ICCs were high for both the new cohort (ICC = 0.94) and the original cohort (ICC = 0.97) ^11^, indicating excellent inter-rater reliability in scoring

### Computational assessments of narrative coherence

Computational linguistic measures were implemented to operationalise two distinct levels of narrative organisation, following discourse coherence frameworks applied to psychiatric and neurological populations ^44,46,127^. Local narrative coherence quantified the semantic relatedness between consecutive sentences within a remembered episodic event, indexing the step-by-step integration of episodic elements. In parallel, global narrative coherence quantified the overall semantic alignment across a remembered event, capturing the stability of the narrative structure. This dual-level analysis enabled the examination of how hippocampal subfield damage differentially affects fine-grained sequential binding versus higher-order narrative organisation during autobiographical memory retrieval.

### Text preprocessing and segmentation

Sentence-BERT (SBERT) is a transformer-based model that generates semantically meaningful sentence embeddings, encoding the contextual meaning of sentences in a continuous vector space. We applied this approach to autobiographical narratives obtained from the Autobiographical Interview to quantify narrative structure in all of the individuals with hippocampal damage and the matched controls.

Prior to computational analysis, all narratives underwent systematic preprocessing. Narratives were segmented into individual sentences using standard rule-based tokenisation (full stops followed by whitespace and capitalisation). To reduce noise from incomplete utterances and filler statements, sentences containing fewer than five tokens were excluded from analysis, which follows established practices in computational discourse analysis ^127^. Each preprocessed event-based narrative was then converted into 384-dimensional semantic embeddings using the all-MiniLM-L6-v2 SBERT model, a transformer-based architecture fine-tuned on semantic textual similarity tasks ^43^.

### Local narrative coherence

Local narrative coherence was operationalised as the mean semantic similarity between adjacent sentences within autobiographical narratives, quantified using cosine similarity between consecutive sentence embeddings. This approach extends established methodologies in computational linguistics and psychiatric research ^44,45,127,128^ to hippocampal amnesia using transformer-based semantic embeddings. Vector transformation through cosine similarity enables direct comparison of sentence embeddings by measuring the angle between vectors in high-dimensional space, thereby quantifying semantic relatedness without requiring explicit linguistic annotation. For a narrative consisting of n sentences with embeddings s₁, s₂, …, sₙ, sequential semantic dependencies were calculated as cosine similarities between each sentence and its immediate predecessor:

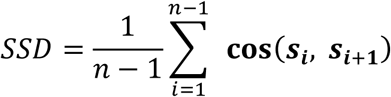

where n denotes the number of sentences in the narrative, sᵢ denotes the embedding vector for the i^th^ sentence, and cos(sᵢ, sᵢ₊₁) denotes cosine similarity between consecutive sentences.

This approach computes the cosine similarity for each adjacent pair of sentences, sums these values across all pairs (from i=1 to i=n-1), and divides by (n-1) to obtain the mean local narrative coherence score.

Local narrative coherence captures the extent to which each sentence semantically builds on the previous sentence to serve as a proxy for the psychological construct of first-order coherence^129,130^, which captures how well consecutive sentences relate to each other.

## Global narrative coherence

Global narrative coherence was quantified as the extent to which individual sentences align with the overarching semantic content of the narrative. Following recent natural language processing approaches ^128,131^, we operationalised global narrative coherence as the mean cosine similarity between each sentence embedding and the centroid of all sentence embeddings within the same narrative:

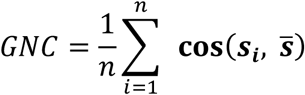

where 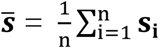 is the mean embedding vector representing the narrative centroid.

This metric captures whether each sentence contributes to a cohesive narrative rather than diverging semantically and corresponds to higher-order coherence constructs in discourse psychology^129,132^. GNC thus quantifies how well individual sentences align with the overall semantic content of the complete remembered episodic event.

### Derived metrics

In order to examine whether group differences in coherence reflected a generalised disruption of narrative structure versus a specific reduction in semantic strength, we extracted distributional properties of the local narrative coherence scores. From the vector of sentence-to-sentence similarities within each narrative, we derived two complementary indices to assess the statistical stability and regularity of semantic transitions, distinct from the mean strength of those transitions. Local narrative coherence variance was calculated as the variance of the cosine similarity scores within each narrative, quantifying the dispersion (from the mean) of semantic connection strengths and indexing the degree of fluctuation in local coherence. This metric assesses the inconsistency of the narrative flow, extending approaches that identified variance in semantic similarity as a discriminative feature in psychiatric populations ^133,134^. Local narrative coherence entropy was calculated as the Shannon entropy of the binned similarity scores (10 bins, range 0–1; i.e., based on a discretised distribution (histogram) of similarity scores), quantifying the distributional uniformity of the transitions and assessing whether the narrative associated with each remembered event sampled a broad range of similarity values (high entropy) or was restricted to a specific range of connection strengths (low entropy).

Together, these metrics differentiate between narratives that are unstable (high variance) versus those that exhibit structural heterogeneity (high entropy), providing a granular examination of how hippocampal integrity contributes to the regulation of narrative quality beyond mean coherence levels ^131,135^.

### Statistical analyses

For all mixed-model ANOVAs, Mauchly’s test assessed the sphericity assumption. When sphericity was violated, Huynh-Feldt corrections were applied to degrees of freedom, with complete sphericity test statistics (χ², p-values, and ε values) reported in Supplementary Information.

Following prior analyses ^11,29,30^, we analysed the composite internal (episodic) and external (semantic) detail scores per participant using omnibus ANOVAs followed by planned comparisons. Given the sample size, we additionally analysed the specific memory response categories (event details, time, place, perceptual details, and emotions/thoughts) that comprise the composite internal and external detail scores. These granular memory content scores were analysed using omnibus ANOVAs followed by planned comparisons, providing complementary detail-level insights.

An alpha criterion of 0.05 was used, with correction for multiple comparisons applied using the Bonferroni-Holm procedure. All *t*-tests, ANOVAs, linear regressions, and Sobel’s test were performed using IBM SPSS Version 29.0 (IBM Corp.).

## Acknowledgements

TDM is supported by the Wellcome Trust (Grant number: 222913/Z/21/Z). AEH is supported by the Medical Research Council (MR/X022013/1 and MR/V007173/1), and the National Institute for Health Research (NIHR) Oxford Health Biomedical Research Centre (BRC). MSZ was supported by National Institute for Health and Care Research University College London Hospitals Biomedical Research Centre. CAA was supported by the National Institute for Health Research (NIHR) Oxford Biomedical Research Centre (BRC) and received research grant support from the UCB-Oxford collaboration, MSD Laboratories, NIHR.

## Author contributions

TDM: conceptualisation, methodology, investigation, data curation, formal analysis, validation, funding acquisition, resources, project administration, visualisation, and writing – original draft, reviewing and editing; JZ: formal analysis and writing – reviewing and editing; AEH: resources, writing – reviewing and editing; TAP: resources, writing – reviewing and editing; PAG: resources, writing – reviewing and editing; CAA: resources and writing – reviewing and editing; MSZ: resources and writing – reviewing and editing; CRR: conceptualisation, methodology, investigation, data curation, formal analysis, validation, funding acquisition, resources, project administration, visualisation, and writing – original draft, reviewing and editing.

For the purpose of Open Access, the author has applied a CC BY public copyright licence to any Author Accepted Manuscript version arising from this submission.

