## Supplementary data for "Sequential narrative binding by hippocampal CA2/3 sustains lifespan episodic detail retrieval"

**Supplementary information**

**Table S1: neuropsychological assessment**

All participants underwent extensive standardised neuropsychological assessment in line with our previous work (detailed in the Methods section).[1,2] Supplementary Table 1 provides the detailed performance of integrated (3.0- and 7.0-Tesla) cohort with hippocampal amnesia compared to the normative Z-transformed data (mean = 0, SD = 0.3), tested using between-group independent-sample two-tailed *t*-tests. The neuropsychological performance for both groups individually has been reported elsewhere.[1,3]

Briefly, the results demonstrated that, compared to the normative data (Supplementary Table 1), the new 3.0-Tesla cohort and the integrated cohort with hippocampal amnesia both exhibited significantly superior performance for verbal and visual intelligence, premorbid intelligence, language, and cognitive flexibility. Only visual memory performance (comprised of the immediate and delayed retrieval of the Rey-Osterreith Complex Figure; highlighted in bold) for the new 3.0-Tesla cohort and integrated cohort were significantly different from normative data, but above the threshold that typically indicates severe impairment ($Z = -1.67$). These data are consistent with selective hippocampal-dependent episodic memory dysfunction rather than global cognitive decline.

**Table S1. Neuropsychological domain performance compared to normative data**

| **Domain** | **n** | **Average Z-score** | **SEM** | **t** | **d.f.** | **p-value** |
|---|---|---|---|---|---|---|
| Verbal intelligence | 32 | 0.96 | 0.11 | 4.07 | 31 | <0.001 |
| Visual intelligence | 32 | 0.83 | 0.08 | 3.51 | 31 | <0.001 |
| Premorbid intelligence | 31 | 0.84 | 0.15 | 3.48 | 30 | <0.001 |
| Verbal memory | 32 | 0.42 | 0.15 | 2.41 | 31 | 0.35 |
| **Visual memory** | **32** | **-0.89** | **0.12** | **-5.12** | **31** | **<0.001** |
| Recognition memory | 32 | -0.16 | 0.13 | -0.92 | 31 | 0.36 |
| Sustained attention | 32 | 0.21 | 0.17 | 1.2 | 31 | 0.24 |
| Language | 32 | 0.79 | 0.18 | 4.54 | 31 | <0.001 |
| Executive function | 32 | 0.28 | 0.1 | 1.58 | 31 | 0.12 |
| Cognitive flexibility | 32 | 0.51 | 0.16 | 2.93 | 31 | 0.006 |
| Visuomotor skills | 32 | 0.22 | 0.09 | 1.28 | 31 | 0.21 |
| Visuoconstruction | 32 | 0.11 | 0.2 | 0.59 | 31 | 0.56 |

**Supplementary data 1: Focal hippocampal subfield damage across both cohorts**

Leucine-rich glioma inactivated 1-limbic encephalitis (LGI1-LE) can lead to hippocampal damage involving cornu Ammonis (CA) 2/3 and CA1 with minimal evidence of extra-hippocampal pathology [4-6], as assessed using high-resolution anatomical MRI, thereby providing a model in which to test causal contributions of CA2/3 and CA1 to episodic consolidation and retrieval. Data from the new 3.0-Tesla cohort with hippocampal amnesia were combined with data from our previously published LGI1-LE cohort [1,2,7-11], yielding an integrated single-aetiology cohort of 32 individuals with CA2/3 and CA1 damage and 32 matched controls, with statistical power for subfield-specific analyses previously accessible only through functional neuroimaging in healthy adults.

In order to harmonise the different segmentation protocols, two separate omnibus ANOVAs were conducted on composite, lateralised dentate gyrus, CA2/3, CA1, and subiculum subfield volumes.

**3.0-Tesla cohort (N = 18 per group)**

A three-way 2 (group: hippocampal amnesia, controls) by 2 (side: left, right) by 4 (subfield: dentate gyrus, CA2/3, CA1, subiculum) mixed-model ANOVA (with Mauchly's test demonstrating that the assumption of sphericity had not been violated) demonstrated that there were significant main effects of the group ($F_{(1,34)} = 14.545, p < 0.001$), side ($F_{(1,34)} = 11.144, p = 0.002$), and subfield ($F_{(3,102)} = 453.774, p < 0.001$). A significant two-way interaction was observed between group and subfield ($F_{(3,102)} = 3.777, p = 0.013$). The three-way interaction was not significant ($F_{(3,102)} = 0.930, p = 0.429$). Planned comparisons demonstrated that the between-group volume loss was isolated to the CA2/3 and CA1 subfields ($F_{(1,34)} = 129.642, p < 0.001, \eta^2 = 0.792$; and $F_{(1,34)} = 15.330, p < 0.001, \eta^2 = 0.311$, respectively; Bonferroni-corrected $p = 0.0125$).

**7.0-Tesla cohort (N = 14 per group)**

A three-way 2 (group: hippocampal amnesia, controls) by 2 (side: left, right) by 4 (subfield: DG, CA2/3, CA1, subiculum) mixed-model ANOVA (with Mauchly's test demonstrating that the assumption of sphericity had been violated for subfield [$\chi^2_{(5)} = 22.542, p < 0.001$] and subfield by side [$\chi^2_{(5)} = 14.483, p = 0.013$], therefore, dfs were corrected using Greenhouse-Geisser estimates for subfield $\varepsilon = 0.64$ and Huynh-Feldt estimates for subfield by side $\varepsilon = 0.77$ respectively) revealed significant main effects of the group ($F_{(1,26)} = 6.314, p = 0.019$), side ($F_{(1,26)} = 8.366, p = 0.008$), and subfield ($F_{(1.927,50.105)} = 71.141, p < 0.001$). A significant two-way interaction was observed between group and subfield ($F_{(1.927,50.105)}$ = 5.010, $p$ = 0.011). The three-way interaction was not significant ($F_{(2.635,68.516)}$ = 0.263, $p$ = 0.827). Planned comparisons demonstrated that this between-group volume loss was isolated to the CA2/3 and CA1 subfields ($F_{(1,26/27)}$ = 10.936, $p$ = 0.003, $\eta^2$ = 0.296; and $F_{(1,26/27)}$ = 8.736, $p$ = 0.008, $\eta^2$ = 0.244, respectively; Bonferroni-corrected $p$ = 0.0125).

**Supplementary data 2: Full statistical analyses of lifespan autobiographical memory retrieval**

The Autobiographical Interview was performed across all participants and underwent transcription for text-based and neurocomputational linguistic analysis. Quantitative assessment revealed no statistical difference between the number of words generated by the integrated (3.0- and 7.0-Telsa) group with hippocampal amnesia (total words: 224793) and the control cohort (total words: 216644 ; $t_{(62)}$ = 0.391, two-tailed $p$ = 0.698, Cohen's d = 0.98).

The new, independent LGI1-LE cohort characterised at 3.0-Tesla replicated findings from the original 7.0-Tesla cohort, revealing extensive anterograde and retrograde amnesia on the Autobiographical Interview (n = 18 participants pre group; N = 36). A 2 (group: amnesic, control) by 2 [memory detail type: internal (episodic), external (semantic)] by 5 [time period: last year (anterograde period); 30-55, 18-30, 11-18, 0-11 (retrograde periods)]. Mauchly's test demonstrated that the assumption of sphericity had been violated for time ($\chi^2_{(9)}$ = 25.495, $p$ = 0.003, ε = 0.898, degrees of freedom (dfs) Huynh-Feldt corrected) and time period by memory detail type ($\chi^2_{(9)}$ = 17.408, $p$ = 0.43, ε = 0.922, dfs Huynh-Feldt corrected). There were significant main effects of group ($F_{(1,34)}$ = 15.702, $p$<0.001, $\eta^2$ = 0.316), time period ($F_{(3.54,122.11)}$ = 10.558, $p$<0.001, $\eta^2$ = 0.237), and memory detail type ($F_{(1,34)}$ = 400.111, $p$<0.001, $\eta^2$ = 0.922). Crucially, there were significant two-way interactions for group by time period ($F_{(3.54,122.11)}$ = 2.585, $p$ = 0.046, $\eta^2$ = 0.071), group by memory detail type ($F_{(1,34)}$ = 11.690, $p$ = 0.002, $\eta^2$ = 0.256), and time by memory detail type ($F_{(3.69,125.42)}$ = 4.246, $p$ = 0.004, $\eta^2$ =0.111). The three-way interaction was not significant ($F_{(3.689,125.416)}$ = 1.811, $p$ = 0.136, $\eta^2$ = 0.051).

A planned comparison for the most remote period (0-11 years) revealed no significant between-group differences for internal (episodic) ($F_{(1,34)}$ = 0.504, $p$ = 0.483, $\eta^2$ = 0.015) nor external (semantic) details ($F_{(1,34)}$ = 2.539, $p$ = 0.116, $\eta^2$ = 0.038). Given these preservations, we conducted a follow-up *post hoc* 2 (group: amnesic group, controls) by 4 [time period: last year (anterograde period); 30-55, 18-30, 11-18 (retrograde periods)] by 2 (memory detail type: internal details, external details) mixed-model ANOVA excluding the 0-11 period. Mauchly's assumption of sphericity had not been violated. Significant main effects of group ($F_{(1,34)}$ = 18.096, $p$<0.001,

$\eta^2 = 0.347$), time ($F_{(3,102)} = 4.437, p = 0.06, \eta^2 = 0.115$), and memory detail type ($F_{(1,34)} = 353.200, p<0.001, \eta^2 = 0.912$) were observed, with significant interaction between group and memory detail type ($F_{(1,34)} = 13.689, p<0.001, \eta^2 = 0.287$) but not group and time period ($F_{(3,102)} = 1.189, p = 0.318, \eta^2 = 0.034$) or memory detail type and time period ($F_{(3,102)} = 2.399, p = 0.072, \eta^2 = 0.066$). The three-way interaction was not significant ($F_{(3,102)} = 0.691, p = 0.559$ $\eta^2 = 0.020$). Planned comparisons on the internal and external detail scores (collapsed over time; alpha criterion Bonferroni-corrected to $p = 0.025$) demonstrated a significant group difference for internal details ($F_{(1,34)} = 17.345, p<0.001, \eta^2 = 0.338$), but not external details ($F_{(1,34)} = 1.315, p = 0.260, \eta^2 = 0.037$), confirming selective episodic memory impairment across four decades.

In the integrated cohort and controls (n = 32 participants per group; N = 64), a 2 (group: amnesic, control) by 2 [memory detail type: internal (episodic), external (semantic)] by 5 [time period: last year (anterograde period); 30-55, 18-30, 11-18, 0-11 years (retrograde periods)] mixed-model ANOVA was conducted on the total internal and external details scores collapsed across the three retrieval cues and response categories constituting the standard administration of the Autobiographical Interview [1,12]. Mauchly's test demonstrated that the assumption of sphericity had been violated for time period ($\chi^2_{(9)} = 35.406, p < 0.001, \varepsilon = 0.849$) and for the time period by memory detail type interaction ($\chi^2_{(9)}= 27.774, p = 0.001, \varepsilon = 0.841$). Degrees of freedom were corrected using Huynh-Feldt estimates. There were significant main effects of group ($F_{(1,62)} = 13.457, p < 0.001, \eta^2 = 0.178$), time period ($F_{(3.68,227.85)} = 7.137, p < 0.001, \eta^2 = 0.103$), and memory detail type ($F_{(1,62)} = 445.496, p < 0.001, \eta^2 = 0.878$). Significant two-way interactions were observed for group by time period ($F_{(3.68,227.85)} = 3.882, p = 0.004, \eta^2 = 0.059$), group by memory detail type ($F_{(1,62)} = 23.067, p < 0.001, \eta^2 = 0.265$), but not time period by memory detail type ($F_{(3.64,225.40)} = 2.279, p = 0.068, \eta^2 = 0.038$). The three-way interaction between group, time period, and memory detail type was not significant ($F_{(3.64,225.40)} = 2.454, p = 0.052, \eta^2 = 0.038$). Planned comparisons for the most remote period (0–11 years) revealed no significant between-group differences for internal (episodic) details ($F_{(1,62)} = 0.504, p = 0.483, \eta^2 = 0.015$) or for external (semantic) details ($F_{(1,62)} = 2.539, p = 0.116, \eta^2 = 0.038$).

Given the sparing of the 0–11 years period, a follow-up 2 (group: amnesic, control) by 2 [memory detail type: internal (episodic), external (semantic)] by 4 [time period: last year (anterograde period); 30–55, 18–30, 11–18 years (retrograde periods)] mixed-model ANOVA was conducted excluding this interval. Mauchly's test indicated that the assumption of sphericity had not been violated. Significant main effects were observed for group ($F_{(1,62)} = 20.822, p < 0.001, \eta^2 = 0.245$), time period ($F_{(3,186)} = 2.972$, p = 0.033, $\eta^2 = 0.044$), and memory detail type

($F_{(1,32)}$ = 408.803, $p$ < 0.001, $\eta^2$ = 0.865). A significant interaction emerged for group by memory detail type ($F_{(1,62)}$ = 25.609, $p$ < 0.001, $\eta^2$ = 0.286). The interactions between group and time period ($F_{(3,186)}$ = 1.366, $p$ = 0.254, $\eta^2$ = 0.021) and between memory detail type and time period ($F_{(3,186)}$ = 0.860, $p$ = 0.463, $\eta^2$ = 0.013) were not significant. The three-way interaction between group, time period, and memory detail type was not significant ($F_{(3,186)}$ = 0.818, $p$ = 0.485, $\eta^2$ = 0.013). Planned comparisons (Bonferroni-corrected $\alpha$ = 0.025) on the internal and external detail scores collapsed over time demonstrated a significant group difference for internal (episodic) details ($F_{(1,62)}$ = 17.345, $p$ < 0.001, $\eta^2$ = 0.338), but not for external (semantic) details ($F_{(1,62)}$ = 1.315, $p$ = 0.260, $\eta^2$ = 0.037), confirming selective episodic memory impairment across anterograde and four retrograde intervals.

In a subset of the integrated cohort, we next assessed the profile of anterograde and retrograde amnesia on a decade-by-decade basis in all participants who were able to generate autobiographical memories from six retrograde decades (N = 36, 18 with amnesia in the subset integrated cohort (3.0-Tesla cohort, n = 8; 7.0-Tesla cohort, n = 10) and 18 matched controls; see Figure 2C–D). A 2 (group: amnesic, control) by 2 [memory detail type: internal (episodic), external (semantic)] by 7 [time period: last year (anterograde period); 50-59, 40–49, 30-39, 20-29, 10-19, 0-9 years (retrograde periods)] mixed-model ANOVA was conducted. Mauchly's test demonstrated that the assumption of sphericity had not been violated. Significant main effects were observed for group ($F_{(1,34)}$ = 32.208, $p$ < 0.001, $\eta^2$ = 0.486), time period ($F_{(6,204)}$ = 2.962, $p$ = 0.009, $\eta^2$ = 0.080), and memory detail type ($F_{(1,34)}$ = 440.115, $p$ < 0.001, $\eta^2$ = 0.924). Significant interactions emerged between group and time period ($F_{(6,204)}$ = 3.705, $p$ = 0.002, $\eta^2$ = 0.098) and between group and memory detail type ($F_{(1,34)}$ = 42.068, $p$ < 0.001, $\eta^2$ = 0.553). The interaction between time period and memory detail type was not significant ($F_{(6,204)}$ = 1.798, $p$ = 0.101, $\eta^2$ = 0.050). The three-way interaction between group, time period, and memory detail type was not significant ($F_{(6,204)}$ = 1.773, $p$ = 0.121, $\eta^2$ = 0.050). Post-hoc comparisons confirmed no significant group differences in internal (episodic) details ($F_{(1,34)}$ = 1.647, $p$ = 0.208, $\eta^2$ = 0.044) or external (semantic) details ($F_{(1,34)}$ = 2.583, $p$ = 0.117, $\eta^2$ = 0.071) for the earliest decade (0-9).

Excluding the preserved earliest decade (0–9 years), a follow-up 2 (group: amnesic, control) x 2 [memory detail type: internal (episodic), external (semantic)] x 6 (time period: last year (anterograde period); 50-59, 40-49, 30-39, 20-29, 10-19 years (retrograde periods)) mixed-model ANOVA was conducted. Mauchly's test demonstrated that the assumption of sphericity had not been violated. Significant main effects were observed for group ($F_{(1,34)}$ = 37.569, $p$ < 0.001, $\eta^2$ = 0.525), time period ($F_{(5,170)}$ = 2.496, $p$ = 0.033, $\eta^2$ = 0.068), and memory detail type ($F_{(1,34)}$ = 414.208, $p$ < 0.001, $\eta^2$ = 0.924). A significant interaction emerged between group and memory detail type ($F_{(1,34)} = 42.716, p < 0.001, \eta^2 = 0.557$). The interactions between group and time period ($F_{(5,170)} = 0.894, p = 0.486, \eta^2 = 0.026$) and time period and memory detail type ($F_{(5,170)} = 1.836, p = 0.108, \eta^2 = 0.051$) were not significant. The three-way interaction between group, time period, and memory detail type was not significant ($F_{(5,170)} = 0.386, p = 0.858, \eta^2 = 0.011$). Planned comparisons (Bonferroni-corrected $\alpha = 0.025$) revealed significant internal (episodic) detail impairments ($F_{(1,34)} = 45.068$, p < 0.001, $\eta^2 = 0.556$) but preserved external (semantic) details ($F_{(1,34)} = 4.175$, p = 0.048, $\eta^2 = 0.104$), demonstrating that hippocampal damage disrupts episodic autobiographical memory across more than 50 years of life. Preservation was limited to early childhood memories, with external (semantic) details spared throughout

**Supplemental data 3: Anterior–posterior transformed CA3 volume *Z*-score and CA1 *Z*-score regression models for internal (episodic) detail scores across time**

Stepwise linear regressions were run with the CA3 and CA1 volume *Z*-scores on the integrated 3.0-Tesla and 7.0-Tesla cohort and controls (see Methods; N = 64). These analyses demonstrated that the transformed anterior CA3 volume *Z*-scores predicted total internal detail scores for all time periods (11-18: $t = 2.108, p = 0.039, R^2 = 0.052, \beta_1 = 0.259$; 18-30: $t = 2.082, p = 0.041, R^2 = 0.050, \beta_1 = 0.256$; 30-55: $t = 3.009, p = 0.004, R^2 = 0.113, \beta_1 = 0.357$; last year: $t = 3.465$, $p<0.001, R^2 = 0.149, \beta_1 = 0.403$), except the 0-11 time period ($t = 0.385, p = 0.721, R^2 = -0.014, \beta_1 = 0.045$). Sobel's test demonstrated that there was a significant indirect effect of group membership mediating the relationship between transformed anterior CA3 *Z*-score and the internal (episodic) detail score (11-18: $Z = 3.196, p = 0.001$, Cohen's d = 0.87; 18-30: $Z = 2.656$, $p = 0.008$, Cohen's d = 0.70; 30-55: $Z = 2.953, p = 0.003$, Cohen's d = 0.79; last year: $Z = 3.987$, $p<0.001$, Cohen's d = 1.15).

Transformed posterior CA3 volumes predicted total internal detail scores for the 18-30 ($t = 2.488, p = 0.016, R^2 = 0.076, \beta_1 = 0.301$), 30-55 ($t = 3.260, p = 0.002, R^2 = 0.146, \beta_1 = 0.383$), and Last Year epochs ($t = 3.278, p = 0.002, R^2 = 0.148, \beta_1 = 0.384$), but not 0-11 ($t = 0.065, p = 0.949, R^2 = -0.016, \beta_1 = 0.008$) or 11-18 ($t = 1.776, p = 0.081, R^2 = 0.033, \beta_1 = 0.220$) time periods. Sobel's test demonstrated that there was a significant indirect effect of group membership mediating the relationship between transformed posterior CA3 *Z*-score and the internal (episodic) detail score (11-18: $Z = 3.349, p = 0.001$, Cohen's d = 0.92; 18-30: $Z = 2.715$, $p = 0.007$, Cohen's d = 0.70; 30-55: $Z = 3.053, p = 0.002$, Cohen's d = 0.83; last Year: $Z = 4.180$, $p<0.001$, Cohen's d = 1.23).

Anterior CA1 *Z*-scores predicted internal (episodic) detail score for the last year ($t$ = 2.325, $p$ = 0.023, $R^2$ < 0.080, $\beta_1$ = 0.283) and no other time period (0-11: $t$ = 0.652, $p$ = 0.517, $R^2$ = 0.007, $\beta_1$= 0.083; 11-18: $t$ = 0.733, $p$ = 0.466, $R^2$ = 0.009, $\beta_1$= 0.093; 18-30: $t$ = 0.949, $p$ = 0.346, $R^2$ = 0.014, $\beta_1$= 0.120; 30-55: $t$ = 1.791, $p$ = 0.078, $R^2$ = 0.049, $\beta_1$= 0.222). Sobel's test demonstrated that there was a significant indirect effect of group membership mediating the relationship between anterior CA1 *Z*-score and the internal (episodic) detail score (11-18: $Z$ = 2.541, $p$ = 0.010, Cohen's d = 0.67; 18-30: $Z$ = 2.251, $p$ = 0.024, Cohen's d = 0.59; 30-55: $Z$ = 2.433, $p$ = 0.015, Cohen's d = 0.64; Last Year: $Z$ = 2.954, $p$ = 0.003, Cohen's d = 1.23).

Posterior CA1 *Z*-scores were similarly found to predict internal (episodic) detail score for the last year ($t$ = 2.799, $p$ = 0.007, $R^2$ = 0.112, $\beta_1$ = 0.335) and no other time period (0-11: $t$ = -1.421, $p$ = 0.160, $R^2$ = 0.032, $\beta_1$= -0.178; 11-18: $t$ = 0.480, $p$ = 0.633, $R^2$ = 0.004, $\beta_1$= 0.061; 18-30: $t$ = 0.699, $p$ = 0.487, $R^2$ = 0.008, $\beta_1$= 0.088; 30-55: $t$ = 1.697, $p$ = 0.054, $R^2$ = 0.059, $\beta_1$= 0.242). Sobel's test demonstrated that there was a significant indirect effect of group membership mediating the relationship between posterior CA1 *Z*-score and the internal (episodic) detail score (11-18: $Z$ = 2.286, $p$ = 0.004, Cohen's d = 0.60; 18-30: $Z$ = 2.463, $p$ = 0.014, Cohen's d = 0.65; 30-55: $Z$ = 2.700, $p$ = 0.007, Cohen's d = 0.72; Last Year: $Z$ = 3.443, $p$ = 0.001, Cohen's d = 0.95).

**Supplementary data 4: Hippocampal damage and context-bound memory components of lifespan episodic retrieval**

In the integrated 3.0-Tesla and 7.0-Tesla cohort and controls (n = 32 participants per group; N = 64), a 2 (group: amnesic, control) by 2 [memory detail type: internal (episodic), external (semantic)] by 5 [time period: last year (anterograde period); 30-55, 18-30, 11-18, and 0-11 years (retrograde periods)] by 5 (memory components: event detail, time, place, perceptual details, emotions/thoughts) mixed-model factorial ANOVA was conducted on the units of information acquired from the standard administration of the AI. Mauchly's test demonstrated that the assumption of sphericity had been violated for time ($\chi^2_{(9)}$ = 34.482, $p$ < 0.001, $\varepsilon$ = 0.93, dfs Huynh-Feldt corrected), memory components ($\chi^2_{(9)}$ = 191.770, $p$ < 0.001, $\varepsilon$ = 0.51, dfs Greenhouse-Geisser corrected), time by memory detail type ($\chi^2_{(9)}$ = 27.878, $p$ < 0.001, $\varepsilon$ = 0.91, dfs Huynh-Feldt corrected), time by memory components ($\chi^2_{(135)}$ = 746.985, $p$ < 0.001, $\varepsilon$ = 0.40, d.f.s Greenhouse-Geisser corrected), memory detail type by memory components ($\chi^2_{(9)}$ = 155.060, $p$ < 0.001, $\varepsilon$ = 0.58, d.f.s Greenhouse-Geisser corrected), time period by memory components by memory detail type ($\chi^2_{(135)} = 675.967$, $p < 0.001$, $\varepsilon = 0.46$, dfs Greenhouse-Geisser corrected).

Main effects of group ($F_{(1,62)} = 16.059$, $p < 0.001$, $\eta^2 = 0.201$), memory detail type ($F_{(1,62)} = 468.313$ $p < 0.001$, $\eta^2 = 0.880$), time period ($F_{(3.68,228.21)} = 7.027$, $p < 0.001$, $\eta^2 = 0.099$), and memory components ($F_{(2.031,125.89)} = 262.671$ $p < 0.001$, $\eta^2 = 0.804$) were found. Two-way interactions between group by time period ($F_{(3.68,228.21)} = 4.523$, $p = 0.002$, $\eta^2 = 0.066$), group by memory detail type ($F_{(1,64)} = 23.067$, $p < 0.001$, $\eta^2 = 0.265$), group by memory components ($F_{(2.031,125.89)} = 4.997$, $p = 0.009$, $\eta^2 = 0.072$), memory components by memory detail type period ($F_{(2.031,125.89)} = 136.575$, $p < 0.001$, $\eta^2 = 0.681$) were all significant, whereas time by memory detail type ($F_{(3.64,225.63)} = 2.305$, $p = 0.070$, $\eta^2 = 0.035$) and time by memory components ($F_{(6.34,392.86)} = 2.305$, $p = 0.065$, $\eta^2 = 0.041$) were not significant. Significant three-way interactions were seen for group, time period, and memory components ($F_{(6.34,392.86)} = 2.769$, $p = 0.032$, $\eta^2 = 0.041$), group, memory components, and memory detail type ($F_{(2.33,144.33)} = 12.574$, $p < 0.001$, $\eta^2 = 0.164$), and time period, memory components, and memory detail type ($F_{(7.299,467.159)} = 3.919$, $p<0.001$, $\eta^2 = 0.058$) but not group, time period, and memory detail type ($F_{(6.346,406.143)} = 2.013$, $p = 0.059$, $\eta^2 = 0.030$). The four-way interaction was also significant ($F_{(7.299,467.159)} = 2.226$, $p = 0.029$, $\eta^2 = 0.034$).

We hypothesised that the internal and external details scores for each of the memory component types would be equivalent across both groups for the most remote period (0-11). Therefore, *post hoc* one-way ANOVAs (Bonferroni corrected $\alpha = 0.005$) revealed that there was no significant between-group differences for internal event ($F_{(1,62)} = 1.043$, $p = 0.311$, $\eta^2 = 0.016$), temporal ($F_{(1,62)} = 1.778$, $p = 0.187$, $\eta^2 = 0.027$), place ($F_{(1,62)} = 0.041$, $p = 0.840$, $\eta^2 = 0.001$), perceptual details ($F_{(1,62)} = 0.036$, $p = 0.849$, $\eta^2 = 0.001$), and emotions/thought details ($F_{(1,62)} = 3.502$, $p = 0.066$, $\eta^2 = 0.052$). The same was true for external event ($F_{(1,64)} = 2.926$, $p = 0.092$, $\eta^2 = 0.044$), temporal ($F_{(1,62)} = 0.189$, $p = 0.665$, $\eta^2 = 0.003$), place ($F_{(1,62)} = 0.136$, $p = 0.189$, $\eta^2 = 0.003$), perceptual ($F_{(1,62)} = 1.805$, $p = 0.184$, $\eta^2 = 0.027$), and emotions/thought details ($F_{(1,62)} = 0.124$, $p = 0.725$, $\eta^2 = 0.070$).

As the planned comparisons demonstrated that there was no significant between-group differences for the most remote period, a further *post hoc* omnibus 2 (group: amnesic, control) by 2 [memory detail type: internal (episodic), external (semantic)] by 4 [time period: last year (anterograde period); 30-55, 18-30, and 11-18 (retrograde periods)] by 5 (memory components: event details, time, place, perceptual details, and emotions/thoughts) mixed-model factorial

ANOVA was conducted with the most remote period removed. Mauchly's test demonstrated that the assumption of sphericity had been violated for time period by memory detail type ($\chi^2_{(77)}$ = 526.963, $p < 0.001$, ε = 0.430, dfs Greenhouse-Geisser corrected), memory detail type by memory component type ($\chi^2_{(9)}$ = 160.192, $p < 0.001$, ε = 0.579, dfs Greenhouse-Geisser corrected), and between time period, memory detail type, and memory components ($\chi^2_{(77)}$ = 465.611, $p < 0.001$, ε = 0.486).

Results revealed significant main effects of group ($F_{(1,62)}$ = 20.822, $p < 0.001$, $\eta^2$ = 0.245), memory detail type ($F_{(1,62)}$ = 265.683 $p < 0.001$, $\eta^2$ = 0.806), time period ($F_{(3,186)}$ = 2.972, $p$ = 0.033, $\eta^2$ = 0.044), and memory components ($F_{(4,248)}$ = 408.803, $p < 0.001$, $\eta^2$ = 0.865). There were significant two-way interactions between group by memory detail type ($F_{(1,62)}$ = 25.609, $p$ = 0.02, $\eta^2$ = 0.286), group and memory components ($F_{(4,248)}$ = 6.709, $p$ = 0.002, $\eta^2$ = 0.095), time period and memory detail type ($F_{(2.762,171.24)}$ = 5.221, $p < 0.001$, $\eta^2$ = 0.075), and memory components and memory detail type ($F_{(2.76,171.24)}$ = 121.569, $p < 0.001$, $\eta^2$ = 0.655), but not group by time ($F_{(3,192)}$ = 1.366, $p$ = 0.254, $\eta^2$ = 0.021) or time period by memory components($F_{(3,192)}$ = 0.860, $p$ = 0.461, $\eta^2$ = 0.013). Significant three-way interactions were seen for group, memory detail type, and memory components ($F_{(2.30,142.52)}$ = 12.677, $p$<0.001, $\eta^2$ = 0.165), and time period, memory detail type, and memory components ($F_{(12,768)}$ = 3.210, $p$<0.001, $\eta^2$ = 0.048), but not between group, time period, and memory detail type ($F_{(12,744)}$ = 1.412, $p$ = 0.155, $\eta^2$ = 0.022), or between group, time period, and memory components ($F_{(12,744)}$ = 0.485, $p$ = 0.155, $\eta^2$ = 0.022). The four-way interaction was also significant ($F_{(12,744)}$ = 2.150, $p$ = 0.012, $\eta^2$ = 0.033).

Given the four-way interaction, *post hoc* omnibus ANOVAs were conducted to assess group by detail type performance across time. Mauchly's test of sphericity was not violated. These ANOVAs demonstrated a significant interaction of group and time for points accrued for temporal details ($F_{(3,186)}$ = 3.907, $p$ = 0.010, $\eta^2$ = 0.058) but not for event details ($F_{(3,186)}$ = 1.381, $p$ = 0.250, $\eta^2$ = 0.021), place ($F_{(3,186)}$ = 0.945, $p$ = 0.420, $\eta^2$ = 0.015), perceptual ($F_{(3,186)}$ = 0.187, $p$ = 0.667, $\eta^2$ = 0.003), and emotions/thoughts details ($F_{(3,186)}$ = 0.705, $p$ = 0.522, $\eta^2$ = 0.011). For each of the memory components, there was a main effect of group: event ($F_{(1,62)}$ = 23.413, $p$<0.001, $\eta^2$ = 0.267), place ($F_{(1,62)}$ = 17.138, $p$<0.001, $\eta^2$ = 0.211), time ($F_{(1,62)}$ = 8.735, $p$ = 0.004, $\eta^2$ = 0.120), perceptual details ($F_{(1,62)}$ = 10.301, $p$ = 0.002, $\eta^2$ = 0.139), and emotions/thought ($F_{(1,62)}$ = 14.286, $p$<0.001, $\eta^2$ = 0.182).

*Post hoc* ANOVAs on the external detail memory components were also conducted. These demonstrated no main effect of group (event: $F_{(1,62)} = 0.408$, $p = 0.525$, $\eta^2 = 0.007$; place: $F_{(1,62)} = 0.156$, $p = 0.694$, $\eta^2 = 0.003$; time: $F_{(1,62)} = 1.210$, $p = 0.276$, $\eta^2 = 0.019$; perceptual details: $F_{(1,62)} = 0.015$, $p = 0.903$, $\eta^2 = 0.000$; and, emotions/thought: $F_{(1,62)} = 0.713$, $p = 0.402$, $\eta^2 = 0.011$) or group by time (event details: $F_{(3,186)} = 2.444$, $p = 0.065$, $\eta^2 = 0.038$; place: $F_{(3,186)} = 1.792$, $p = 0.150$, $\eta^2 = 0.028$; time: $F_{(3,186)} = 1.653$, $p = 0.179$, $\eta^2 = 0.026$; perceptual ($F_{(3,186)} = 0.770$, $p = 0.512$, $\eta^2 = 0.012$; and, emotions/thoughts details: ($F_{(3,186)} = 0.882$, $p = 0.451$, $\eta^2 = 0.014$).

**Supplementary data 5: Long-axis hippocampal subfield volume *Z*-scores and episodic memory detail type regression models**

To examine structure–function relationships across hippocampal subfields, we conducted regression model-based analyses and mediation analyses using total CA2/3, CA3 and CA1 volumes as predictors, with the total internal (episodic) detail and memory components scores from the Autobiographical Interview as dependent measures.

In the integrated 3.0- and 7.0-Tesla cohort and controls (N = 64), linear regression analyses revealed that total transformed CA3 volume *Z*-scores significantly predicted each context-bound component comprising the internal (episodic) detail score (excluding the 0-11 period internal detail subcategories): event ($t = 4.256$, $p<0.001$, $R^2 = 0.226$, $\beta_1 = 0.476$), place ($t = 2.565$, $p = 0.013$, $R^2 = 0.096$, $\beta_1 = 0.310$), time ($t = 2.777$, $p = 0.007$, $R^2 = 0.096$, $\beta_1 = 0.333$), perceptual ($t = 2.481$, $p = 0.016$, $R^2 = 0.090$, $\beta_1 = 0.300$), emotion/thought details ($t = 3.346$, $p<0.001$, $R^2 = 0.162$, $\beta_1 = 0.403$) and total internal (episodic) details ($t = 4.242$, $p<0.001$, $R^2 = 0.212$, $\beta_1 = 0.474$). For CA3 volume z-score, mediation results using Sobel's test demonstrated significant indirect effects of group membership on these relationships: total internal (episodic) details (Z = 4.499, p < 0.001, Cohen's d = 1.36), event (Z = 4.425, p < 0.001, d = 1.33), place (Z = 3.879, p < 0.001, d = 1.11), time (Z = 2.702, p < 0.001, d = 0.72), perceptual (Z = 2.948, p = 0.003, d = 0.79), and emotion/thought details (Z = 3.438, p < 0.001, d = 0.95).

When excluding the most remote period, transformed CA2/3 volume *Z*-scores predicted total internal (episodic) detail performance across the lifetime ($t = 3.780$, $p<0.001$, $R^2 = 0.187$, $\beta_1 = 0.433$), total event details ($t = 3.817$, $p<0.001$, $R^2 = 0.190$, $\beta_1 = 0.436$), total place detail ($t = 2.014$, $p = 0.048$, $R^2 = 0.046$, $\beta_1 = 0.248$), total temporal details ($t = 2.465$, $p = 0.016$, $R^2 = 0.089$, $\beta_1 = 0.299$), total perceptual details ($t = 2.208$, $p = 0.031$, $R^2 = 0.073$, $\beta_1 = 0.270$), and total emotion/thought details ($t = 3.250$, $p = 0.002$, $R^2 = 0.146$, $\beta_1 = 0.381$).

For the CA2/3 volume *Z*-scores, there was an indirect effect of group membership on the relationship between total CA2/3 *Z*-score and memory component from the Autobiographical Interview: total internal details (Sobel's *Z*-score = 4.186, $p<0.001$, Cohen's d = 1.23); total event details (Sobel's $Z = 3.627$, $p<0.001$, Cohen's d = 1.02); total place score (Sobel's $Z = 3.387$, $p = 0.001$, Cohen's d = 0.94); total temporal details (Sobel's $Z = 2.508$, $p = 0.012$, Cohen's d = 0.66); total perceptual details (Sobel's $Z = 2.705$, $p = 0.007$, Cohen's d = 0.72); and total emotions/thoughts details (Sobel's $Z = 3.212$, $p = 0.001$, Cohen's d = 0.88).

By contrast, total CA1 volume predicted only total internal detail ($t = 2.084$, $p = 0.041$, $R^2 = 0.065$, $\beta_1 = 0.256$) and event detail performance ($t = 2.210$, $p = 0.031$, $R^2 = 0.73$, $\beta_1 = 0.270$), with non-significant associations found for total place details ($t = 0.952$, $p = 0.345$), temporal details ($t = 1.645$, $p = 0.105$), perceptual details ($t = 1.044$, $p = 0.300$), or emotion/thought details ($t = 1.965$, $p = 0.054$). Mediation analyses revealed significant indirect effects of group membership on the relationship between transformed CA1 *Z*-score and both total internal ($Z = 3.402$, $p = 0.002$, Cohen's d = 0.425) and event detail scores ($Z = 3.395$, $p = 0.001$, Cohen's d = 0.424).

**Supplementary data 6: Hippocampal damage impairs local narrative coherence but not global narrative coherence and local narrative variance and entropy**

In the integrated 3.0- and 7.0-Tesla cohort and controls (N = 64), a 2 (group: amnesic, control) by 5 [time period: last year (anterograde period); 30-55, 18-30, 11-18, 0-11 (retrograde periods)] mixed-model ANOVA was conducted on local narrative coherence cosine similarity scores for each memory. Mauchly's test demonstrated that the assumption of sphericity had been violated for time ($\chi^2_{(9)} = 24.985$, $p = 0.003$, $\varepsilon = 0.812$, dfs Huynh-Feldt corrected). There was a main effect of group ($F_{(1,62)} = 4.251$, $p = 0.043$, $\eta^2 = 0.064$) but not time period ($F_{(3.51,217.38)} = 0.835$, $p = 0.504$, $\eta^2 = 0.013$). There was no significant two-way interaction between group and time period ($F_{(3.15,217.38)} = 1.817$, $p = 0.141$, $\eta^2 = 0.028$).

Given the absence of between-group differences in AI detail scores for the most remote period (see Supplementary data 1–2), we examined whether local narrative coherence showed a similar pattern. A follow-up one-way between-group ANOVA on the local narrative coherence cosine similarity scores for the most remote period demonstrated no significant between-group difference ($F_{(1,62)} = 0.493$, $p = 0.485$, $\eta^2 = 0.008$). We then performed a follow-up *post hoc* 2 (group: amnesic, control) by 4 [time period: last year (anterograde period); 30-55, 18-30, 11-18, (retrograde periods)] mixed-model ANOVA on local narrative coherence cosine similarity scores for each memory. Mauchly's test demonstrated that the assumption of sphericity had been violated for time ($\chi^2_{(9)}$ = 18.882, $p$ = 0.002, ε = 0.816, dfs Huynh-Feldt corrected). There was a main effect of group ($F_{(1,62)}$ = 6.824, $p$ = 0.011, $\eta^2$ = 0.099) but not time period ($F_{(2.596,160.974)}$ = 0.834, $p$ = 0.463, $\eta^2$ = 0.013). There was no significant two-way interaction between group and time period ($F_{(2.596,160.974)}$ = 0.857, $p$ = 0.451, $\eta^2$ = 0.014).

Next, the global narrative coherence cosine similarity scores for each memory were analysed 2 (group: amnesic, control) by 5 (time period: last year (anterograde period); 30-55, 18-30, 11-18, 0-11 years (retrograde periods)] mixed-model ANOVA. Mauchly's test demonstrated that the assumption of sphericity had been violated for time ($\chi^2_{(9)}$ = 32.073, $p$<0.001, ε = 0.778, dfs Huynh-Feldt corrected. There was no significant main effect of group ($F_{(1,62)}$ = 0.790, $p$ = 0.378, $\eta^2$ = 0.013) but a significant main effect was seen for time period ($F_{(3.346,207.463)}$ = 4.352, $p$ = 0.004, $\eta^2$ = 0.066). There was no significant two-way interaction between group and time period ($F_{(3.346,207.463)}$ = 1.797, $p$ = 0.142, $\eta^2$ = 0.028).

An omnibus 2 (group: amnesic, control) x 5 [time period: last year (anterograde period); 30-55, 18-30, 11-18, 0-11 (retrograde periods)] mixed-model ANOVA was conducted on the local narrative coherence variance scores generated from each memory. Mauchly's test demonstrated that the assumption of sphericity had not been violated. There was no significant main effect of time period ($F_{(4,348)}$ = 0.437, $p$ = 0.782, $\eta^2$ = 0.007) nor group ($F_{(1,62)}$ = 0.009, $p$ = 0.925, $\eta^2$ = 0.000). The two-way interaction between group and time period was not significant ($F_{(4,248)}$ = 1.559, $p$ = 0.186, $\eta^2$ = 0.025).

An omnibus 2 (group: amnesic, control) x 5 [time period: last year (anterograde period); 30-55, 18-30, 11-18, 0-11 years (retrograde periods)] mixed-model ANOVA was conducted on the local narrative coherence entropy values generated from each memory. Mauchly's test demonstrated that the assumption of sphericity had not been violated. There was no significant main effect of time period ($F_{(4,248)}$ = 1.080, $p$ = 0.367, $\eta^2$ = 0.017) nor group ($F_{(1,62)}$ = 0.234, $p$ = 0.630, $\eta^2$ = 0.004). The two-way interaction between group and time period was also not significant ($F_{(4,248)}$ = 1.166, $p$ = 0.326, $\eta^2$ = 0.018).
